# An Endocytic Checkpoint Controls Macrophage PD-1 Function and Immunotherapy Fate

**DOI:** 10.64898/2026.04.14.718292

**Authors:** Madhubanti Mullick, Ella McLaren, Suchismita Roy, Brandon Biagas, Mahitha Shree Anandachar, Vanessa Castillo, Samuel Williams, Celia R. Espinoza, Courtney Tindle, Gajanan D. Katkar, Patricia A. Thistlethwaite, Saptarshi Sinha, Pradipta Ghosh

## Abstract

Responses to PD1 blockade span durable tumor control to hyperprogressive disease (HPD), yet innate immune mechanisms governing these extremes remain undefined. Here we integrate a macrophage systems atlas (>12,500 transcriptomes) with single-cell profiles from >1,000 anti–PD1 treated patients, to identify *CCDC88A* (**GIV**) as a macrophage-intrinsic determinant of durable response versus HPD. GIV loss increases PD1 surface retention, suppresses phagocytosis, and accelerates tumor growth across murine models, human macrophages, and patient-derived organoids. Myeloid-specific GIV deletion converts PD1 blockade from tumor-restraining to tumor-accelerating by reprogramming macrophages toward HPD-like states. Mechanistically, GIV engages a conserved TIR-like [TILL] motif within the PD1 cytoplasmic tail to drive dynamin-dependent endocytosis, coupling innate immune signaling logic to checkpoint receptor trafficking. Pharmacologic disruption of this axis phenocopies GIV loss, revealing an endocytic vulnerability that undermines checkpoint efficacy and triggers accelerated growth at relapse. These findings define PD1 routing, rather than ligand-binding, as a macrophage-encoded checkpoint governing antitumor immunity.

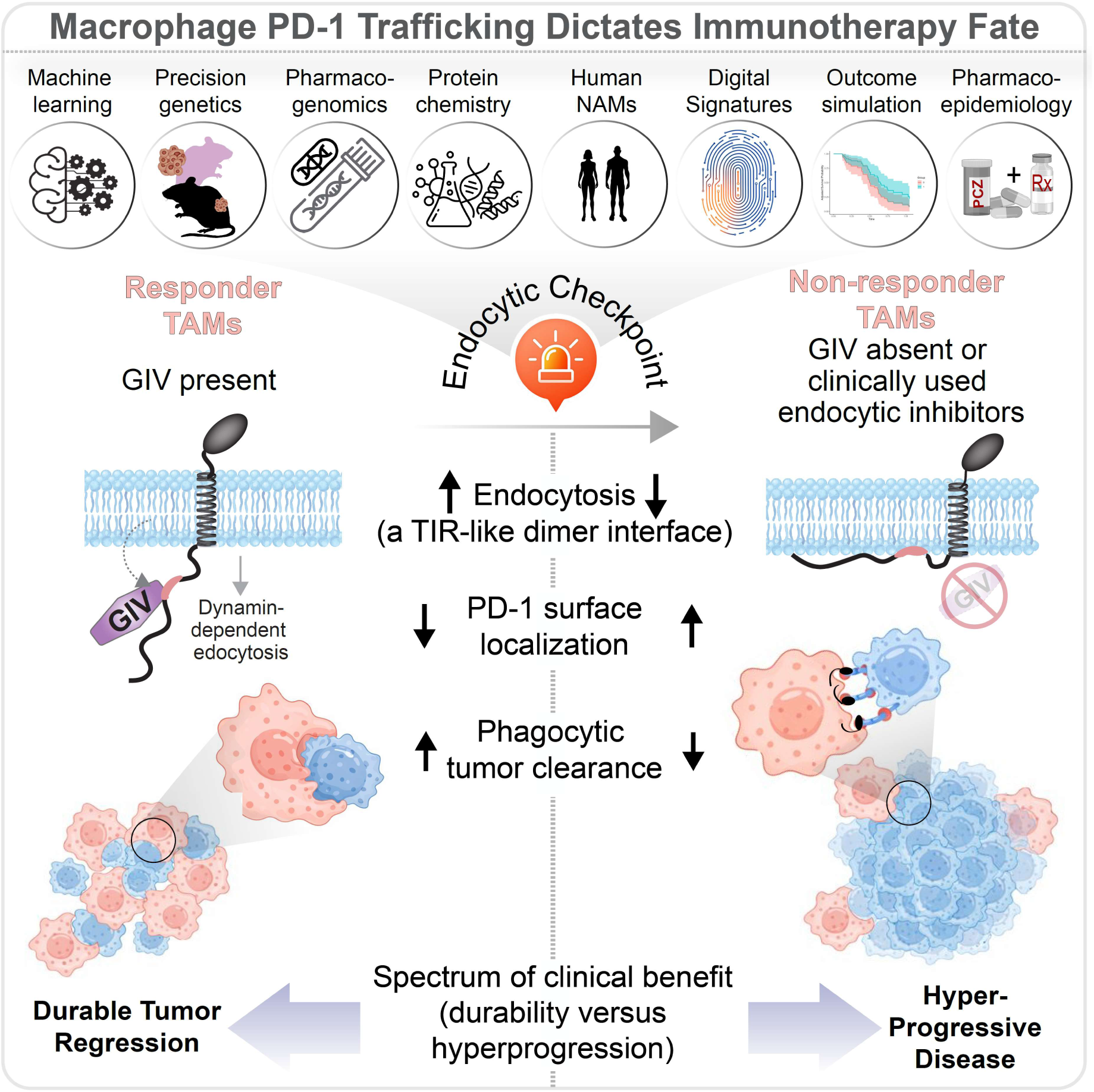
GRAPHIC ABSTRACT.

**In brief:** Mullick *et al.* show that PD1 trafficking in tumor associated macrophages (TAMs), controlled by the endocytic adaptor GIV/Girdin, determines the durability of immunotherapy. Disrupting this pathway traps PD1 at the surface, converts checkpoint blockade into hyperprogressive disease, and abolishes therapeutic benefit, revealing PD1 routing as a macrophage-encoded checkpoint of immunotherapy outcome.

**Highlights:**

- GIV defines macrophage states that stratify durable response vs hyperprogressive disease
- GIV limits surface checkpoint accumulation by facilitating PD1 endocytosis
- Loss of GIV traps PD1 at the membrane and drives hyperprogressive disease
- Endocytic blockade phenocopies GIV loss and abolishes checkpoint efficacy
- Psychotropic drugs disrupt PD1 trafficking and accelerate tumor growth

## INTRODUCTION

Immune checkpoint inhibitors (**ICI**s) have reshaped cancer therapy, delivering durable tumor regression in a subset of patients across diverse malignancies^1^. Yet clinical responses remain strikingly heterogeneous. While some patients achieve long-term durable remission, others fail to respond or, more alarmingly, develop secondary resistance and progress despite treatment. When progression is accompanied by accelerated tumor growth after ICI exposure, the phenomenon is termed hyperprogressive disease (**HPD**)--a paradoxical acceleration of tumor growth rate following therapy^2^. HPD occurs in ∼6% to 43% of patients^3^ and is defined by a 50%^4^ to >2-fold^5^ increase in tumor growth rate after treatment relative to baseline. It occurs more frequent when treated with single-agent checkpoint blockade and is associated with high metastatic burden and poor prognosis^4^. Despite its clinical impact, the biological mechanisms and actionable biomarkers that distinguish durable benefit from accelerated progression remain poorly defined^6–8^ (**Figure 1A**).

**Figure 1:**
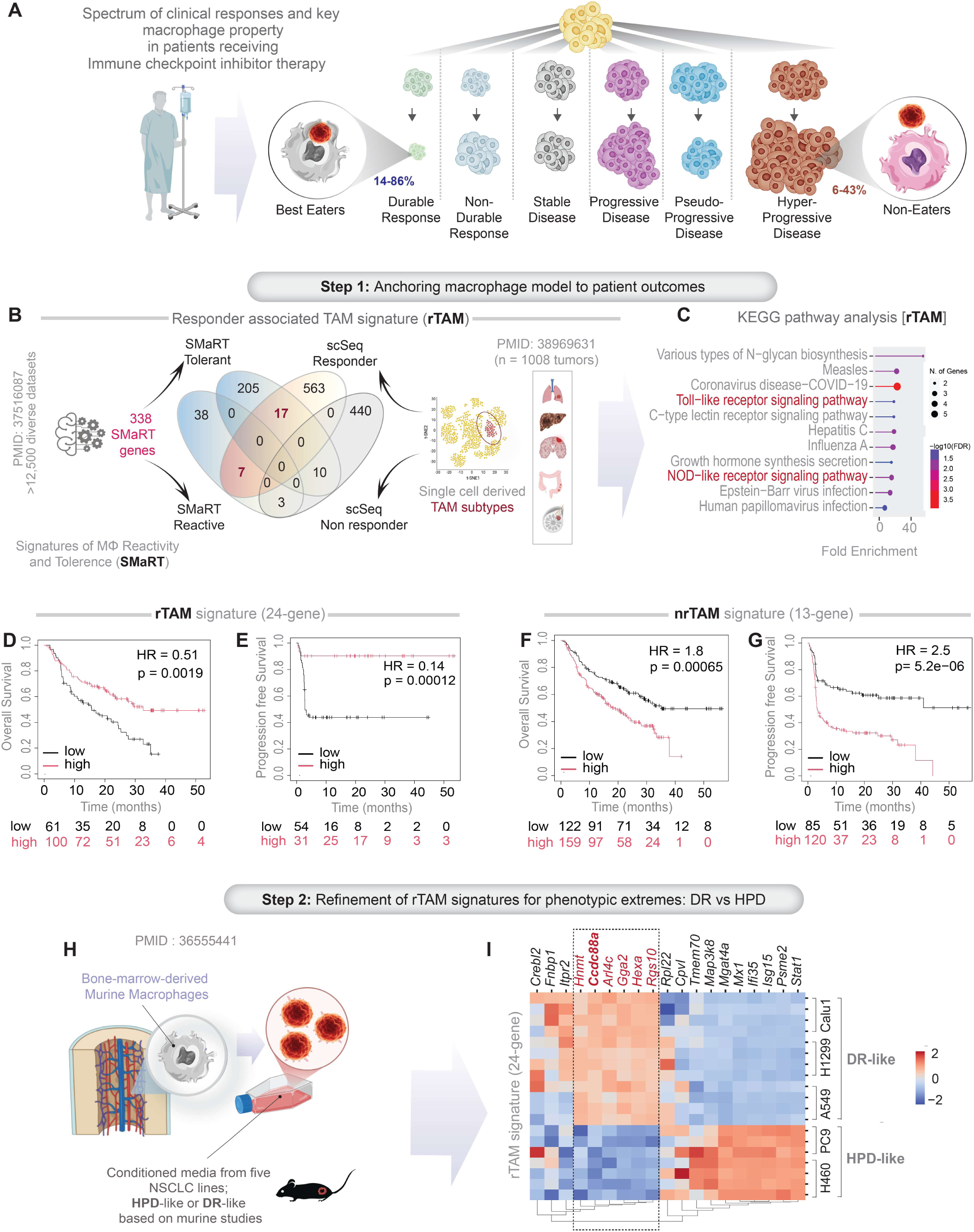
Systems anchoring of macrophage states to extremes of immunotherapy response. (**A**) Schematic depicting the clinical spectrum of outcomes to immune checkpoint inhibitor (ICI) therapy, ranging from durable response to hyperprogressive disease (HPD). % reflects estimated frequencies across solid tumors^99–101^. Macrophage effector capacity (“best eaters” vs “non-eaters”) is highlighted as a candidate determinant investigated in this study. (**B-I**) Integrative 2-step computational framework to identify tumor-associated macrophage (TAM) programs linked to immunotherapy outcomes. **Step 1:** Anchoring macrophage states to patient outcomes (**B-G**). A machine learning-derived model of macrophage continuum process—Signatures of Macrophage Reactivity and Tolerance (SMaRT)^45^—was integrated with single-cell TAM profiles from anti–PD-1–treated patient tumors. A Venn diagram **(B)** shows overlap between SMaRT, derived from >12,500 transcriptomes (n = 771 human, 1,651 murine), and single-cell TAM clusters (n=363,315 TAMs or macrophage-like cells) from the CPI1000+ anti–PD1 cohort (n = 1,008 patients)^46^. Overlapping genes define responder-associated (rTAM; 24 genes) and non-responder-associated (nrTAM; 13 genes) signatures. Full gene lists are provided in **Data S1**. KEGG pathway enrichment (C) of rTAM genes, highlighting innate immune and pattern-recognition receptor signaling pathways (red). See also **Figure S1A-F** for additional GO and gene-level analyses. Kaplan–Meier analyses (**D-G**) demonstrating prognostic performance of rTAM (D, E) and non-responder TAM (nrTAM; F, G) signatures for overall and progression-free survival in anti–PD1–treated patients (high versus low expression) across 9 different cancers. **Step 2** (**H-I**): Refinement toward phenotypic extremes (**H-I**). Experimental workflow (H) to refine rTAM signatures using bone marrow–derived macrophages exposed to conditioned media from NSCLC cell lines exhibiting DR-like or HPD-like growth in vivo^48^. Heatmap (I) shows unsupervised clustering of rTAM genes based on z-normalized expression pattern across five NSCLC models identifies subsets associated with durable response versus hyperprogression. Full gene lists are provided in **Data S2**. *Statistics*: Survival analyses used log-rank tests.

Programmed cell death protein 1 (**PD1; CD279**) is a central inhibitory immune checkpoint that regulates T cell function^9,10,11,12^ and maintains immune homeostasis in various physiological states, including infection and autoimmunity^13,14^. Engagement of PD1 by its ligands PD-L1 (CD274) and PD-L2 (CD273) enforces peripheral tolerance and also suppresses anti-tumor immunity^15,16^. Therapeutic antibodies targeting the PD1–PD-L1 axis have revolutionized oncology^17,18,19^. However, as with other ICIs, only a fraction of patients derive durable benefit, and clinical outcomes have largely plateaued despite combination strategies. This has fueled renewed interest in understanding how PD1 transmits inhibitory signals at the molecular level, beyond ligand binding alone.

Although **PD1** blockade was originally framed as a T cell–centric intervention^20–24^, this paradigm cannot fully explain resistance, relapse, or HPD^25^. PD1 is also expressed in innate immune cells^26–30^, including tumor-associated macrophages (**TAMs**), which are increasingly recognized as upstream determinants of tumor progression and immunotherapy fate^31–36^ and potential drivers of HPD^37^. PD1 expression on TAMs inversely correlates with proinflammatory polarization and phagocytic capacity^26,38^. Furthermore, genetic studies^39^ have implicated myeloid-PD1, rather than T cell PD1, as a critical regulator of anti-tumor immunity.

Emerging clinical and experimental evidence from T cell studies indicates that PD1 endocytosis, rather than surface blockade *per se*, is required to unleash cytolytic immune activity^40^. Bivalent antibodies that promote receptor clustering and internalization (e.g., nivolumab) outperform monovalent antibodies (e.g., pembrolizumab) or PD-L1 targeting antibodies^40–43^. While this paradigm has been explored in T cells, the role of PD1 trafficking in macrophages, particularly its surface retention and endocytic control, remains largely unexplored. This is despite the known ability of endocytic inhibitors to enhance phagocytosis and antibody-dependent cellular cytotoxicity (ADCC)^44^ for immunoglobulin (IgG)1-based biologics.

Here, we reframe durable response and the emergence of secondary resistance presenting as HPD as biological extremes to interrogate the full spectrum of TAM programs that govern immunotherapy outcome. By integrating systems immunology, single-cell tumor profiling, and functional models, we identify *CCDC88A* (GIV/Girdin) as a macrophage-intrinsic endocytic adaptor that controls PD1 surface availability. Our findings establish PD1 trafficking—not ligand engagement alone—as a macrophage-encoded checkpoint that determines therapeutic success or failure. Moreover, we reveal a clinically relevant vulnerability: commonly used drugs that inhibit endocytosis may impose a biphasic risk by transiently enhancing checkpoint engagement while ultimately undermining durability through sustained retention of PD1 on the macrophage surface.

## RESULTS

### Identification of macrophage states associated with extremes of checkpoint response

We sought to define TAM programs that govern response to PD-1 blockade. Rather than contrasting average responders with non-responders, we hypothesized that the mechanistic logic of immunotherapy fate would be most discernible at clinical extremes. We therefore deployed a two-step, outcome-extreme discovery strategy (**Figure 1B-I**) designed to resolve macrophage states that actively enable durable benefit versus those that drive therapeutic failure.

In Step 1, we overlaid our machine-learning–derived macrophage continuum model—the Signatures of Macrophage Reactivity and Tolerance (SMaRT)^45^, constructed from >12,500 transcriptomes—onto single-cell tumor datasets from the CPI1000+ cohort (>1,000 anti–PD-1–treated patients)^46^. The SMaRT model organizes macrophage behavior along the full continuum between the extremes of reactivity and tolerance^47^. This cross-scale alignment enabled outcome-anchored mapping of macrophage states rather than lineage abundance. Intersection analyses revealed discrete gene programs enriched within responder versus non-responder TAM clusters, defining responder-associated (rTAM) and non-responder (nrTAM) signatures (**Figure 1B**; **Figure S1A**; **Data S1**).

Pathway enrichment analyses demonstrated that rTAMs programs are enriched for innate immune and pathogen-sensing programs, including TLR and NOD-like receptor signaling, as well as interferon-mediated antiviral responses and heterotrimeric G protein–related molecular functions (**Figure 1C**; **Figure S1B–C**). In contrast, nrTAMs were dominated by ribosomal biogenesis, translational control, and RNA-binding functions, consistent with metabolically active but immunologically inert macrophages (**Figure S1D–F**). Clinically, high rTAM expression robustly predicted improved overall and progression-free survival following PD1 blockade, whereas nrTAM enrichment predicted poor outcomes (**Figure 1D–G**), validating the outcome relevance of this systems framework.

In Step 2, we refined these programs using an experimental extreme-phenotype model. We leveraged a study^48^ that transcriptionally profiled macrophage education *in vitro* using conditioned media from non-small cell lung cancer (NSCLC) cell lines that display either durable response–like (DR-like) or HPD–like growth in mice (**Figure 1H**). Unsupervised clustering of rTAM genes across these polarized conditions resolved discrete modules segregating DR-associated from HPD-associated macrophage states (**Figure 1I**). By intentionally leveraging phenotypic extremes—clinical and experimental—this strategy amplifies biological signal over noise, enabling resolution of macrophage decision circuits that would be obscured in intermediate states.

Together, this outcome-extreme systems approach defines a macrophage effector axis that tracks immunotherapy fate and establishes a conceptual and analytical framework to uncover macrophage-encoded checkpoints governing response durability versus hyperprogression.

### *CCDC88A* (GIV) couples the responder macrophage state to PD1 surface control

Among rTAM genes, *CCDC88A* (encoding **GIV**; Gα-interacting vesicle-associated protein, also known as Girdin) emerged as the sole candidate that simultaneously (i) tracks with durable clinical response in patients, (ii) integrates signals downstream of innate immune receptors, including multiple TLRs (including TLR4^49^) and NOD-like receptors (i.e., NOD2^50^), and (iii) directly modulates heterotrimeric Gα protein signaling^51–56^. As the prototypical non-receptor *<u>G</u>*uanine nucleotide *<u>E</u>*xchange *<u>M</u>*odulator (GEM^51^), GIV links diverse receptor classes to G protein signaling^57^—unlike canonical GPCR-restricted signaling^49,58,59^—and is highly expressed in macrophages^49^, where it is a key determinant of polarization within the SMaRT framework^60^. These properties nominate GIV as a putative mechanistic bridge between macrophage state, innate immune signaling, and PD1 pathway outcomes.

To test this hypothesis *in vivo*, we used mice with myeloid-specific deletion of *CCDC88A*^49,61^ (see *Methods*) to create a syngeneic LL/2 lung cancer model (**Figure 2A**). Loss of GIV resulted in significantly increased tumor burden, reflected by elevated tumor volume and weight at harvest (**Figure 2A**). Tumors from GIV-KO mice exhibited altered distribution of PD1-positive immune cells, restricted to tumor periphery (**Figure 2B**). Critically, a macrophage-intrinsic increase in PD1 surface expression on CD11b⁺F4/80⁺ TAMs was observed by flow cytometry (**Figure 2C**). Overall immune cell composition within the tumor microenvironment was unchanged (**Figure S2B–C**), demonstrating that GIV specifically regulates PD1 in myeloid cells.

**Figure 2:**
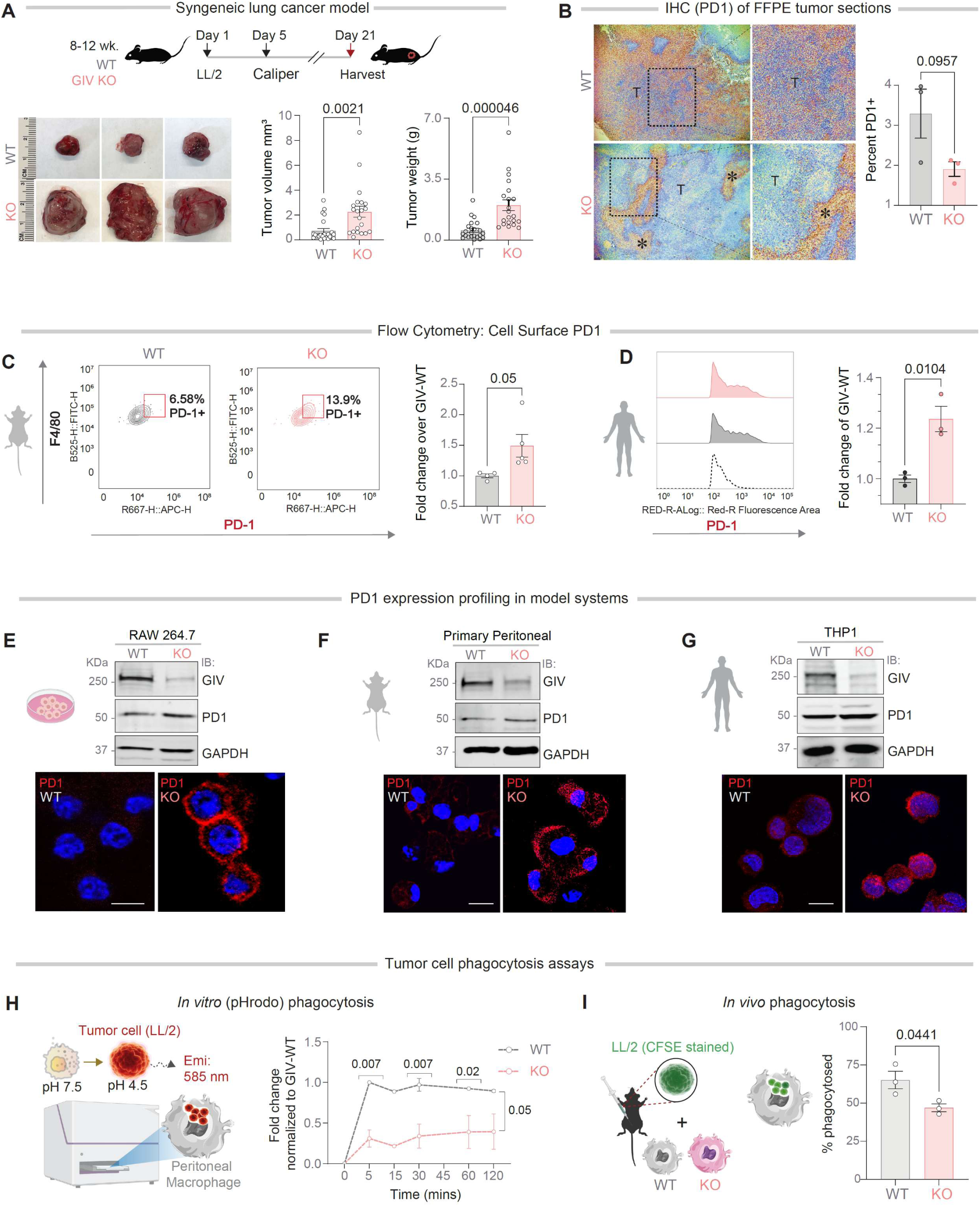
*CCDC88A* deletion increases PD1 on macrophage surface and impairs tumor phagocytosis. **(A)** *In vivo* syngeneic lung cancer model. Schematic (top) of experimental design using WT and myeloid-specific *CCDC88A* (GIV)–deficient (GIV-KO) mice subcutaneously implanted with Lewis lung carcinoma (LL/2) cells. Representative tumors (bottom-*left*) and quantification of tumor volume (bottom-*middle*; mm^3^) and weight (bottom-*middle*; g) at harvest are shown. **(B)** Immunohistochemical (IHC) staining of PD1 in FFPE tumor sections from WT and GIV-KO mice (left), with quantification of percent PD1–positive area (right). **(C)** Flow cytometric analysis of surface PD1 on tumor-infiltrating macrophages from enzymatically digested tumors. Representative contour plots (left) and quantification of PD1 mean fluorescence intensity (MFI) are shown (right). Gating strategy is provided in **Figure S2A**. **(D)** Flow cytometric histograms (left) and quantification (right) of surface PD1 on PMA-differentiated THP1 macrophages following GIV depletion. See **Figure S2B-C** for other immune cell profiles. **(E–G)** PD1 expression across macrophage model systems. Immunoblotting (top) and immunofluorescence staining (bottom) of RAW 264.7 murine macrophages (E), thioglycolate-elicited primary peritoneal macrophages **(H)** *In vitro* phagocytosis assay. Schematic of the pHrodo-based phagocytosis assay (left) and time-course quantification of LL/2 lung tumor cell uptake by murine peritoneal macrophages (right). See also **Figure S3A-C** for similar studies using human MDA-MB-231 breast tumor lines and THP1-derived macrophages. **(I)** *In vivo* phagocytosis assay. Experimental schematic depicting uptake of CFSE-labeled LL/2 cells following intraperitoneal injection (left) and quantification of phagocytosed tumor cells within F4/80⁺ macrophages (right). *Statistics:* Data are shown as mean ± SEM (n = 3–5 biological replicates). Statistical significance was assessed using unpaired t test and Tukey’s multiple comparison test (for *in vitro* phagocytosis assay) as indicated. *p*-values < 0.05 were considered significant.

To establish robustness and cross-species reproducibility, we validated this phenotype across *three* independent macrophage model systems: (i) GIV-depleted RAW 264.7 murine macrophages (∼85–90% knockdown; **Figure 2E**); (ii) thioglycolate-elicited primary peritoneal macrophages from myeloid-specific GIV knockout mice (∼85–90% depletion; **Figure 2F**); and (iii) human THP1 macrophages with CRISPR-mediated GIV loss (>90% depletion; **Figure 2G**). Across all systems, GIV depletion consistently increased macrophage surface PD1, as assessed by flow cytometry, immunoblotting, and immunofluorescence (**Figure 2D–G**), demonstrating a conserved and macrophage-intrinsic role for GIV in restraining PD1 surface accumulation.

Together, these data establish GIV as an essential regulator of surface PD1 levels on macrophages across *in vivo*, primary, and immortalized cell-line models. By linking responder macrophage state to a concrete receptor-localization phenotype, GIV emerges as a decisive rTAM-intrinsic determinant of antitumor immunity and sets the stage for mechanistic dissection of how PD1 trafficking governs immunotherapy response.

### Macrophage GIV licenses phagocytosis and tumor clearance in human organoid models

Because macrophage PD1 functions as a “don’t-eat-me” checkpoint^26^, its accumulation on the cell surface is predicted to suppress phagocytosis. Consistent with this, loss of GIV impaired macrophage phagocytic function: GIV-deficient primary peritoneal macrophages showed markedly reduced tumor cell uptake *in vitro* (**Figure 2H**; **Figure S3A–C**). Similarly, GIV-deficient macrophages (F4/80⁺) engulfed fewer fluorescently (CFSE)-labeled tumor cells *in vivo* (**Figure 2I**).

We next asked whether this defect translates into impaired clearance of patient-derived tumors in a human-relevant system. To address this, we established a human macrophage–tumor organoid co-culture assay in which CFSE-labeled patient-derived lung adenocarcinoma (LUAD) organoids (PDOs) were incubated with WT and GIV-KO THP1-derived macrophages (**Figure 3A**). PDOs were rigorously benchmarked against matched donor tissue to confirm preservation of tumor identity, architecture, and lineage markers (TTF1^62,63^, CK7)^64^ (**Figure S3D–F**). To preserve continuity with other functional readouts and prioritize genetic interpretability over patient-specific immune heterogeneity, we used THP1 macrophages rather than donor-matched immune cells. GIV loss resulted in significantly increased residual tumoroid burden after 4 h of co-culture, reflected by right-shifted size-distribution histograms and increased residual tumoroid area (**Figure 3C–E**). By contrast, adjacent normal lung organoids were cleared equivalently by WT and GIV-KO macrophages, with no significant difference in residual organoid area (**Figure 3E**). Because PDO generation efficiency, histopathologic integrity, and marker expression were indistinguishable across co-culture conditions (**Figure S3D–F**), the observed differences in tumor clearance are attributable to macrophage genotype rather than tumor-intrinsic variability.

**Figure 3:**
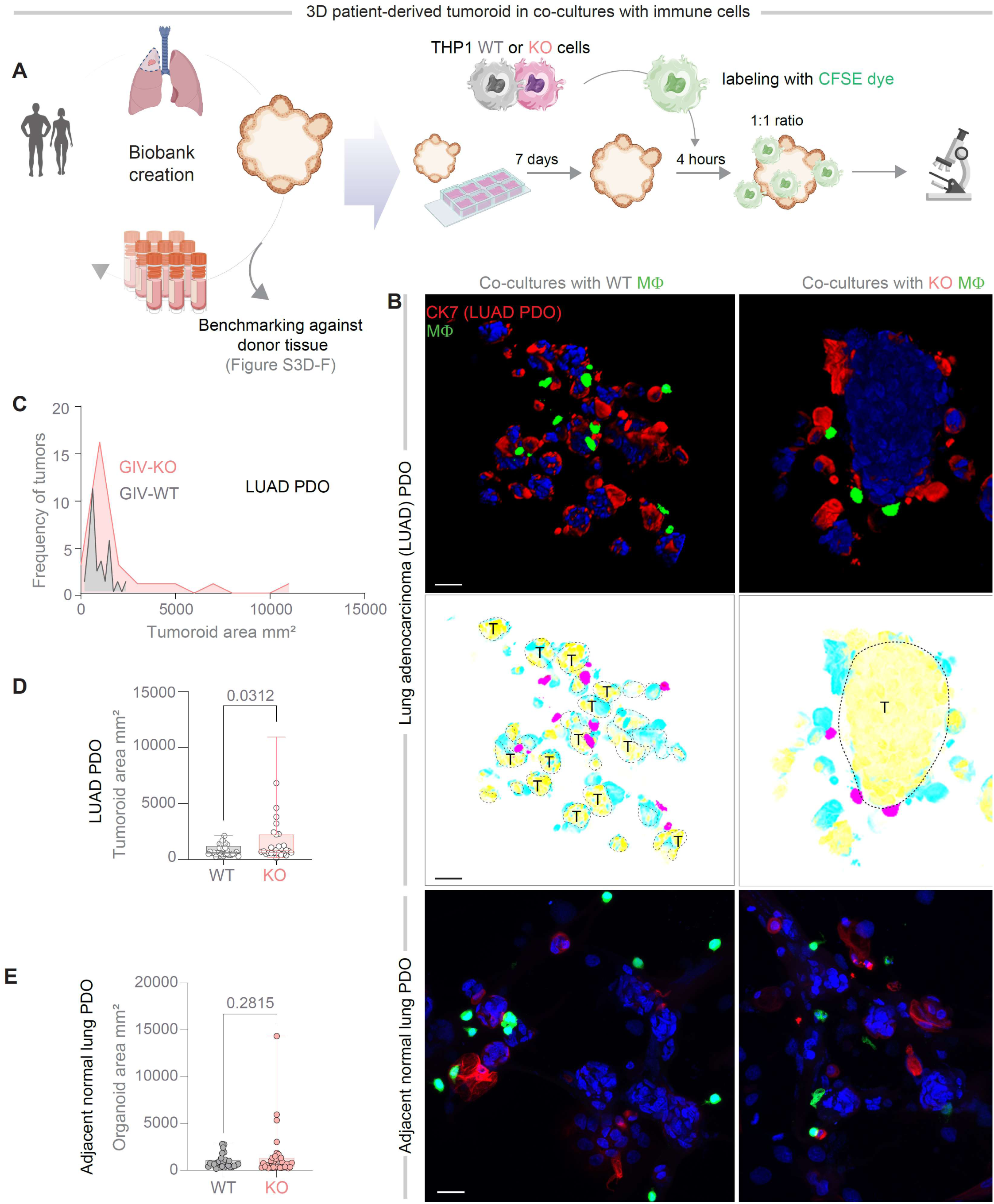
Macrophage GIV (*CCDC88A*) governs tumor organoid clearance in a human co-culture model. **(A)** Schematic of a human co-culture assay in which patient-derived lung tumor organoids (PDOs) are incubated with CFSE-labeled, PMA-differentiated THP1 macrophages that are either WT or GIV-deficient. PDOs were established from lung adenocarcinoma specimens, benchmarked against matched donor tissue (see **Figure S3D–F**), and co-cultured with macrophages at a 1:1 ratio for 4 h. Patient demographics are provided in **Table 1**. **(B)** Representative confocal images and 3D reconstructions of PDOs following 4 h co-culture with THP1 macrophages. Top, lung adenocarcinoma (LUAD) PDOs, shows as merged fluorescent channels and with pseudocolored overlays highlighting tumor boundaries and macrophage positioning; bottom, adjacent normal lung PDOs. Scale bar = 100 µm. **(C-E)** Quantification of residual organoid area after 4 h co-culture with WT versus GIV-KO THP1 macrophages, shown as size-distribution histograms of tumoroid area (C), whisker plots of residual LUAD PDO area (D), and whisker plots of residual adjacent normal lung organoid area (E). *Statistics:* Data are shown as mean ± SEM (n = 2 biological, and 2 technical replicates). Statistical significance was assessed using unpaired t test. *p*-values < 0.05 were considered significant.

These findings were in keeping with high-resolution time-lapse imaging in reductionist 2D assays, where CFSE-labeled MDA-MB-231 breast cancer cells were incubated with WT and GIV-KO THP1-derived macrophages on fibronectin-coated plates (**Figure S3A**). WT macrophages underwent rapid tumor cell engulfment, often completing phagocytosis after a single productive encounter, whereas tumor cells survived multiple failed interactions with GIV-deficient macrophages (**Figure S3B-C**).

Together, findings across three orthogonal systems — reductionist 2D assays, *in vivo* phagocytosis, and human tumor organoid co-culture systems — establish GIV as a macrophage-intrinsic checkpoint for tumor clearance. The restriction of this phenotype to malignant, but not normal, organoids underscores that GIV-dependent macrophage function is context-aware and tumor-specific, positioning GIV as a functional executor of TAM-mediated antitumor immunity.

### Myeloid GIV favors PD1 blockade–induced durable response; its loss unleashes hyper progression

We next asked whether loss of macrophage GIV alters the therapeutic trajectory of PD1 blockade *in vivo*. We treated tumor-bearing WT and myeloid-specific GIV-KO mice with anti–PD1 in a syngeneic LL/2 lung cancer model (**Figure 4A**). At the end of treatment (week 3), both genotypes appeared to respond: anti–PD1 slowed tumor growth and reduced tumor burden relative to untreated controls (**Figure 4B-C**; AUC analyses in **Figure S4A–B**), despite the higher baseline tumor burden in GIV-KO mice (**Figure 4B-C**). Thus, initial tumor shrinkage alone suggested ‘success’ in both genotypes.

**Figure 4:**
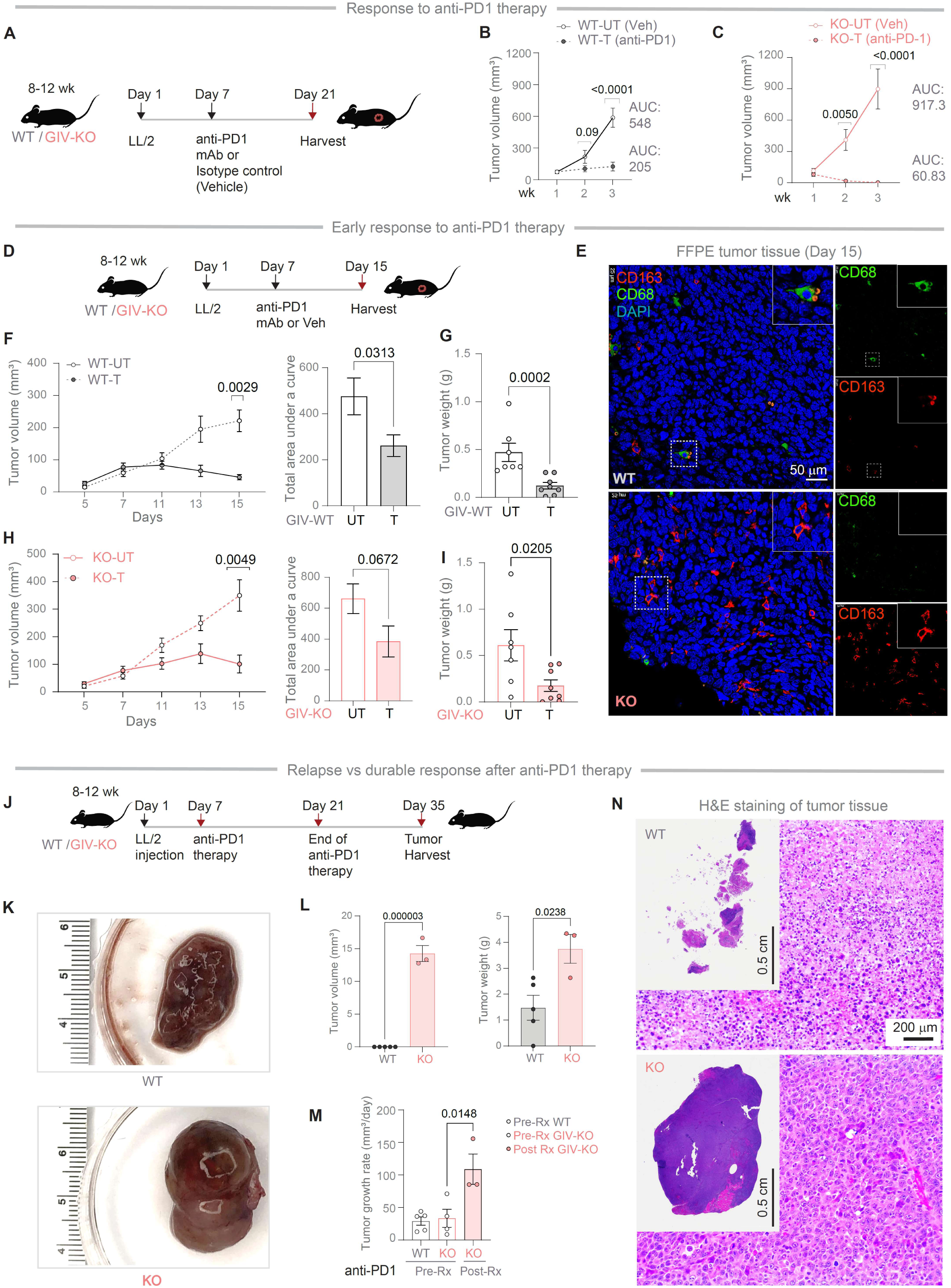
Myeloid GIV loss converts PD1 blockade into accelerated tumor growth *in vivo*. **(A)** Experimental schematic of anti–PD1 administration in a syngeneic ectopic LL/2 lung tumor model using WT and myeloid-specific CCDC88A (GIV)–deficient (LysM-Cre) mice. **(B, C)** Tumor growth kinetics following anti–PD1 therapy in WT (B) and GIV-KO (C) mice compared with untreated controls, revealing tumor control in both WT and GIV-KO mice. Area-under-the-curve (AUC) analyses are shown in **Figure S4A–B**. **(D)** Schematic of the early-response paradigm with tumor harvest at day 15 after therapy initiation (See **Figure S4C** for tumor images). **(E)** Immunofluorescence of FFPE tumor sections showing macrophage infiltration patterns (CD68, green; CD163, red; DAPI, blue). WT tumors show macrophage accumulation within the tumor core, whereas GIV-KO tumors exhibit peripheral enrichment. **(F, G)** Early tumor response in WT mice, assessed by longitudinal tumor volume measurements and AUC (F) and confirmed by reduced tumor weight at harvest (G). **(H, I)** Parallel analyses in GIV-KO mice demonstrate similar early therapeutic benefit, confirmed by volume measurements and AUC (H) and tumor weight (I). **(J)** Experimental timeline to assess relapse versus durable response after cessation of anti–PD1 therapy. **(K)** Representative post-treatment tumors at relapse. WT tumors exhibit liquefied tumor beds with prominent fibrous encapsulation, whereas GIV-KO tumors retain solid tumor architecture. **(L)** Quantification of tumor burden at relapse. Tumor volume (left) reflects measurable solid tumor mass only, whereas tumor weight (right) includes fibrous capsule and exudative material. **(M)** Tumor growth rates before (Pre-Rx) and after (Post-Rx) anti-PD1 therapy. **(N)** Representative H&E-stained tumor sections corresponding to samples in (K), illustrating liquefied, acellular tumor beds in WT mice versus densely cellular tumors in GIV-KO mice. *Statistics:* Data are mean ± SEM (n = 3–10 biological replicates). Statistical significance was determined by Tukey’s multiple comparisons test (tumor growth represented as Area under a curve (AUC)), Bonferroni’s multiple comparisons for tumor growth and response to anti-PD1 therapy, unpaired t test for AUC and harvested tumor weights, volumes and growth rates, as indicated. *p*-values < 0.05 were considered significant.

Because early response is a poor surrogate for durable benefit^65^ — a meaningful and the most desirable immunotherapy clinical trial endpoint^66,67^— we explicitly separated initial sensitivity from long-term outcome. In an early-response paradigm with harvest at day 15 (**Figure 4D**; **Figure S4C**), WT and GIV-KO mice showed comparable early responses, with similar reductions in tumor volume and weight (**Figure 4F–I**; **Figure S4D**). These data establish that macrophage GIV and GIV-dependent surface control of PD1 expression are dispensable for the initial response to PD1 blockade.

Despite similar early responses, tumor architecture and macrophage organization diverged sharply. WT tumors displayed deep macrophage penetration into the tumor core, whereas GIV-KO tumors showed peripheral macrophage confinement, a spatial pattern associated with ineffective clearance (**Figure 4E**). This divergence predicted radically different fates. After treatment cessation (**Figure 4J**), WT mice maintained durable tumor control, marked by liquefied tumor beds encased in fibrotic capsules—hallmarks of immune-mediated clearance (**Figure 4K**). In stark contrast, GIV-KO mice underwent rapid tumor rebound, developing large, solid tumors consistent with hyperprogressive disease (**Figure 4K**). Relapse analyses confirmed significantly increased tumor volume and weight in GIV-KO mice (**Figure 4L**), with a ∼2-3-fold accelerated tumor growth rate (**Figure 4M**) and histopathology revealing high tumor cellularity and persistent malignant glandular architecture (**Figure 4N**). Notably, in WT mice achieving durable response, tumor beds became liquefied and exudative (**Figure 4K**). This phenotype precluded accurate caliper-based volume measurements and rendered post-harvest weight an unreliable surrogate for viable tumor burden (**Figure 4L, 4N**)

Taken together, these results identify myeloid GIV as a gatekeeper of PD1 blockade durability. Dispensable for initial response, but essential for sustained tumor control, GIV prevents the emergence of secondary resistance and accelerated progression at relapse; its loss converts PD1 blockade from a tumor-restraining therapy into a tumor-accelerating liability by disabling macrophage PD1 regulation and phagocytic clearance.

### GIV preserves immune and endocytic transcriptional programs that forecast durable PD1 blockade

We next asked whether GIV shapes tumor and macrophage transcriptional programs associated with durable response to PD1 blockade. Using the syngeneic LL/2 model, we profiled both whole tumors and magnetically enriched F4/80⁺ TAMs from WT and GIV-KO mice following anti–PD1 therapy (**Figure 5A–B**). Loss of myeloid GIV induced broad transcriptional rewiring of the tumor microenvironment (**Figure 5C**). Genes enriched in WT tumors mapped to innate immune, Fc receptor, and inflammatory signaling pathways, whereas these programs were selectively lost in GIV-KO tumors (**Figure 5D**). Importantly, WT-enriched tumor gene signatures robustly predicted improved overall and progression-free survival in patients treated with PD1/PD-L1 (**Figure 5E**), but not CTLA-4 (**Figure S5**)–targeting ICIs, establishing pathway specificity for GIV-dependent programs.

**Figure 5:**
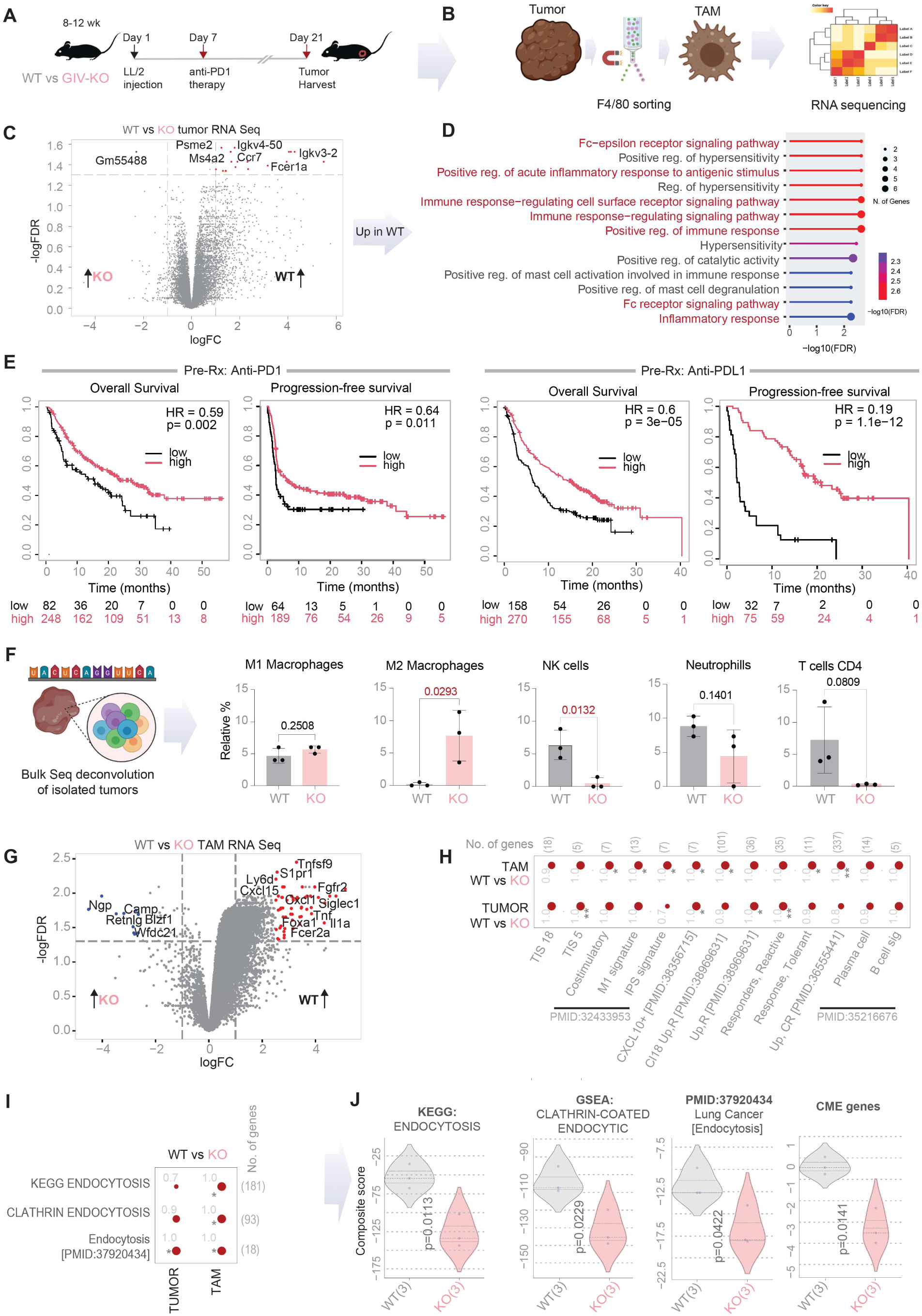
Myeloid GIV preserves innate immune signaling and endocytic programs that predict durable response to PD1 blockade. **(A-B)** Experimental and analytical workflow. *(A)* Schematic of a syngeneic ectopic LL/2 lung tumor model treated with anti–PD1 in WT and GIV-KO mice. *(B)* Tumors were enzymatically dissociated, F4/80⁺ TAMs were magnetically enriched, and *both* whole tumors and TAMs were subjected to RNA sequencing. **(C)** Volcano plot of differentially expressed genes in whole tumors (WT vs. GIV-KO; Adj p-value ≤ 0.05, |logFC| ≥1), highlighting broad transcriptional rewiring upon myeloid GIV loss. Full gene lists are provided in **Data S3**. **(D)** KEGG pathway enrichment of genes upregulated in WT tumors, revealing preservation of innate immune, Fc receptor, and inflammatory signaling programs. **(E)** Kaplan–Meier analyses showing that WT-enriched tumor gene signatures predict improved overall and progression-free survival in patients treated with the class of ICI therapeutics, across 9 different cancers, targeting the PD1•PDL1 axis, i.e., anti–PD1 (left) or anti–PD-L1 (right) ICI (see **Figure S5** for anti–CTLA-4). **(F)** *In silico* deconvolution of bulk tumor transcriptomes showing shifts in immune cell composition, with selective loss of effector macrophage and NK cell signatures in GIV-KO tumors. See Figure 2D and **Figure S2B-C** for experimentally determined immune cell profiles. **(G)** Volcano plot of differentially expressed genes in F4/80⁺ TAMs (WT vs. GIV-KO), identifying macrophage-intrinsic transcriptional reprogramming. Full gene lists are provided in **Data S4**. **(H)** Dot plots comparing enrichment of immune and macrophage state signatures in TAMs versus whole tumors; dot size reflects classification accuracy (AUC) and color denotes directionality (red, upregulated; blue, downregulated). List of genes in each signature is provided in **Data S5**. **(I)** Dot plots, as in H, comparing endocytosis related gene signatures. **(J)** Violin plots compare composite scores for endocytic and clathrin-mediated trafficking gene signatures in F4/80⁺ TAMs from WT v GIV-KO tumors. *Statistics*: Differential expression used adjusted p-value, calculated using Benjamini-Hochberg correction. Survival analyses used log-rank tests. Statistical significance for signature comparisons was calculated using Welch’s t-test and annotated using standard codes (*p ≤ 0.05; **p ≤ 0.01; ***p ≤ 0.001). *p*-values < 0.05 were considered significant.

*In silico* deconvolution of bulk tumor transcriptomes revealed a selective depletion of effector macrophage and NK cell gene signatures in GIV-KO tumors (**Figure 5F**), consistent with impaired immune function despite otherwise preserved immune cell abundance in the tumor microenvironment (**Figure 2D**; **Figure S2B–C**). Focusing on macrophage-intrinsic effects, transcriptomic profiling of F4/80⁺ TAMs uncovered extensive reprogramming upon GIV loss (**Figure 5G**). Cross-cohort and cross-species benchmarking using a rigorous panel of multiple human-relevant gene signatures of ICI-response revealed that WT TAMs were enriched for reactive responder-associated TAM states, whereas GIV-KO TAMs shifted toward tolerant, non-responder–associated TAM programs (**Figure 5H**). Strikingly, among the most selectively depleted programs in GIV-KO TAMs were endocytosis and clathrin-mediated trafficking pathways (**Figure 5I–J**), implicating loss of receptor trafficking capacity as a core feature of GIV deficiency.

These insights into macrophage-intrinsic transcriptional state suggest that GIV-proficient cells retain intact innate immune signaling and endocytic competence—programs that selectively predict durable response to PD1/PD-L1 blockade^40,68,69^. In contrast, GIV loss dismantles these pathways, uncoupling checkpoint inhibition from sustained antitumor immunity. Importantly, this state-level reprogramming is consistent with GIV’s established function as a dynamin-binding scaffold^51^ and gatekeeper of selective clathrin-mediated endocytosis (CME) of receptors^69–73^. Together, these findings nominate GIV as a macrophage-intrinsic molecular checkpoint that links transcriptional state to clinical ICI outcome—driving response when present and resistance when absent.

### GIV is an endocytic adaptor that routes PD1 and determines immunotherapy durability

We hypothesized that GIV may dictate the divergent outcomes of PD1 blockade by physically coupling to and controlling CME of PD1. Co-immunoprecipitation studies (**Figure 6A**) and *in situ* proximity ligation assays (PLA^74^; **Figure 6B**) independently confirmed that endogenous full-length PD1 and GIV proteins formed complexes in THP1-derived macrophages. Inhibiting dynamin-dependent endocytosis with the highly specific, low-toxicity inhibitor DynGo4a^75^ further increased PD1•GIV complexes (**Figure 6B**; **Figure S6A–B**), consistent with accumulation at the plasma membrane and arrested clathrin-mediated endocytic intermediates. Orthogonal validation in a heterologous system showed enhanced peripheral PD1 retention selectively in GIV-KO—but not WT—HeLa cells (**Figure S6C**), establishing a direct, GIV-dependent effect on PD1 surface residence.

**Figure 6:**
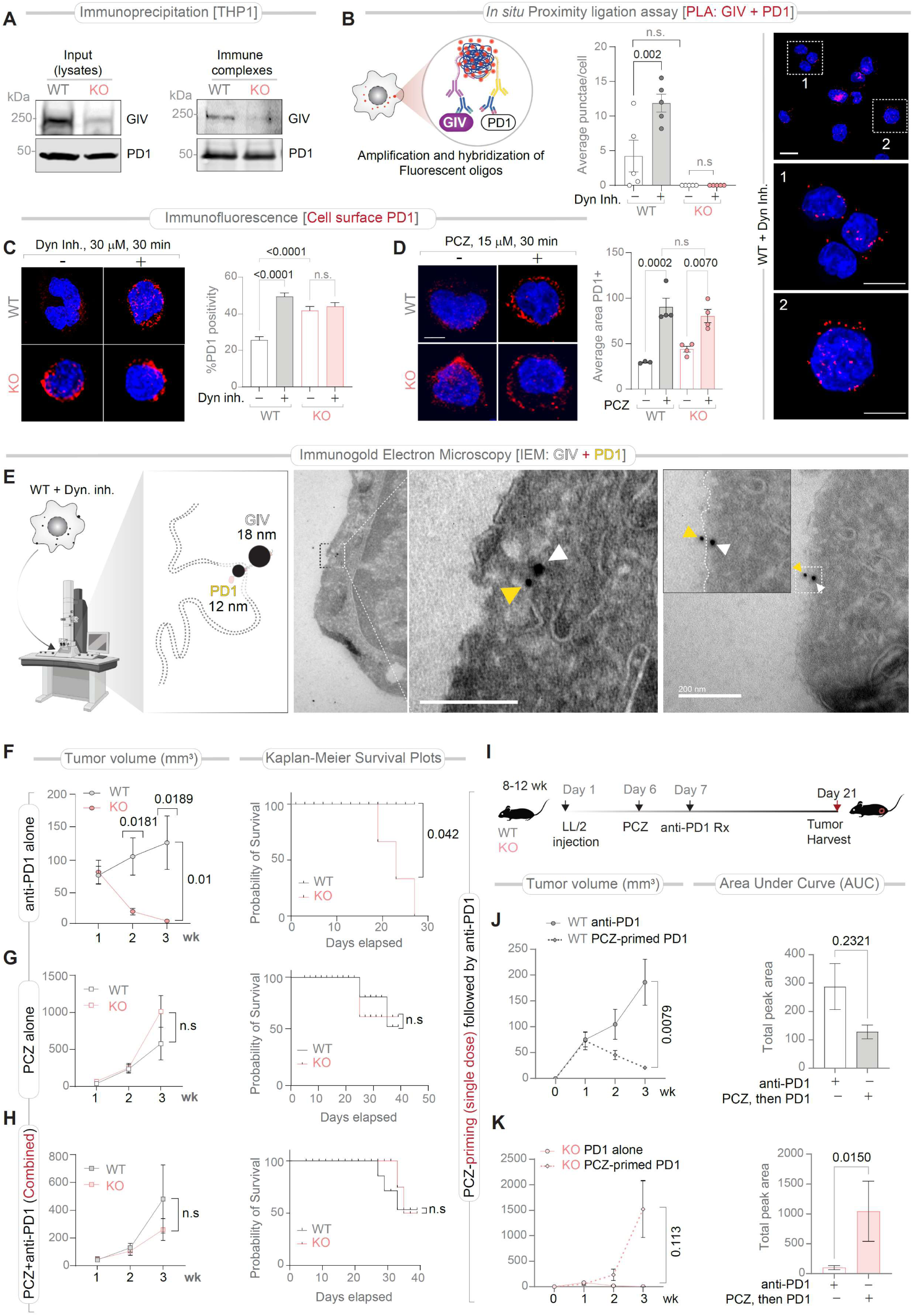
GIV-dependent PD1 endocytic routing determines durability of anti–PD1 immunotherapy. **(A)** Co-immunoprecipitation of endogenous PD1 with GIV in THP1 macrophages from WT and GIV-KO cells. Immunoblots show GIV and PD1 in input lysates and immune complexes. **(B)** *In situ* proximity ligation assay (PLA) detecting endogenous GIV•PD1 interactions. Schematic (left), representative confocal images (right), and quantification (middle) show reduced GIV•PD1 proximity upon inhibition of dynamin-dependent endocytosis (DynGo4a). **(C, D)** Confocal imaging and quantification of surface PD1 in WT versus GIV-KO THP1 macrophages following inhibition of endocytosis with DynGo4a (C) or the prescription drug prochlorperazine (PCZ) (D), revealing exaggerated PD1 surface retention in the absence of GIV. See **Figure S6A-B** for additional fields and controls. **(E)** Immunogold electron microscopy (IEM) demonstrating nanoscale co-localization of GIV (18 nm gold) and PD1 (12 nm gold) at sites of PM invagination following acute dynamin inhibition. See **Figure S6C** for orthogonal immunofluorescence-based localization studies in HeLa cells. **(F–H)** *In vivo* therapeutic responses. Tumor growth and Kaplan–Meier survival analyses in WT and myeloid-specific GIV-KO mice treated with anti–PD1 alone (F), PCZ alone (G), or anti–PD1 plus PCZ (administered concurrently; H). **(I)** Experimental schematic for PCZ priming followed by anti–PD1 therapy. **(J, K)** Tumor growth kinetics and AUC quantification show that PCZ priming enhances anti–PD1 efficacy in WT mice (J) but fails—and exacerbates tumor growth—in GIV-KO mice (K). *Statistics:* Data are mean ± SEM (n = 3–10 biological replicates). Statistical significance was assessed by one-way ANOVA with Bonferroni correction for PLA quantification, Bonferroni multiple comparisons test for tumor growth response to treatment, unpaired t-test for AUC, and Log-rank (Mantel-Cox) test for Kaplan–Meier survival analyses. *p*-values < 0.05 were considered significant.

Functionally, blocking endocytosis unmasked a strict requirement for GIV in PD1 surface control. In WT macrophages, DynGo4a or the clinically used inhibitor prochlorperazine (**PCZ**) modestly increased surface PD1 (**Figure 6C–D**). In contrast, these perturbations produced no additional effect beyond the already exaggerated PD1 surface accumulation in GIV-KO macrophages—i.e., pharmacologic inhibition phenocopied genetic GIV loss (**Figure 6C–D**). Immunogold electron microscopy localized PD1 and GIV to sites of plasma membrane invagination following acute dynamin inhibition, placing the complex at the endocytic interface (**Figure 6E**).

We then tested therapeutic consequences *in vivo*. Anti–PD1 restrained tumor growth and improved survival in WT mice but failed in GIV-KO mice (**Figure 6F**). PCZ alone worsened outcomes in WT mice (**Figure 6G**), erasing genotype-dependent differences and phenocopying GIV loss (**Figure 2A**). Strikingly, concurrent PCZ administration abrogated anti–PD1 efficacy in WT mice, driving trajectories indistinguishable from GIV-KO animals (**Figure 6H**), demonstrating that sustained endocytic blockade nullifies PD1 therapy. Aggressive tumor growth in these PCZ-treated cohorts precluded longer term analyses of relapse/durability of response because all mice reached IACUC mandated endpoints for euthanasia.

Because transient trafficking perturbations (with drugs like PCZ) may differ from sustained inhibition (genetic deletion of GIV), we tested a priming strategy, administering a single dose of PCZ 24 h before the initiation of anti–PD1 (**Figure 6I**), guided by studies documenting a short plasma half-life and longer tumor accumulation^76^. Brief PCZ priming enhanced anti–PD1 efficacy in WT mice, reducing tumor growth and AUC relative to anti–PD1 alone (**Figure 6J**). By contrast, the same regimen failed in GIV-KO mice (**Figure 6K**), consistent with compounding an intrinsic endocytic defect.

These findings demonstrate that GIV facilitates CME of PD1 away from the macrophage surface to sustain antitumor immunity. Disrupting this trafficking—genetically or pharmacologically—traps PD1 at the surface, compromises macrophage phagocytic function, and converts PD1 blockade from durable therapy into failure. Thus, the duration and context of endocytic interference dictate PD1 efficacy, placing receptor routing at the core of immunotherapy durability. Notably, because PCZ is routinely used as an anti-emetic agent in cancer care and proposed as an adjuvant with ADCC-mediated antibodies^77,44,78,79^ these findings carry immediate translational caution and relevance.

### A conserved TILL motif couples GIV to PD1 to control receptor trafficking

We next asked how GIV physically engages PD1 to control receptor routing. Domain mapping revealed that GIV’s C-terminus and the PD1 cytoplasmic tail each harbor a short, conserved TIR-like loop (TILL) motif (**Figure 7A–B**), originally described in cytokine receptors^80^ and shared with TIR-containing innate immune receptors^49^. Because TILL-motifs mediate homotypic interactions^49^, this convergence in sequence suggested that PD1 co-opts an evolutionarily conserved innate-immune interaction logic, i.e., a putative TILL-motif to engage GIV.

**Figure 7.**
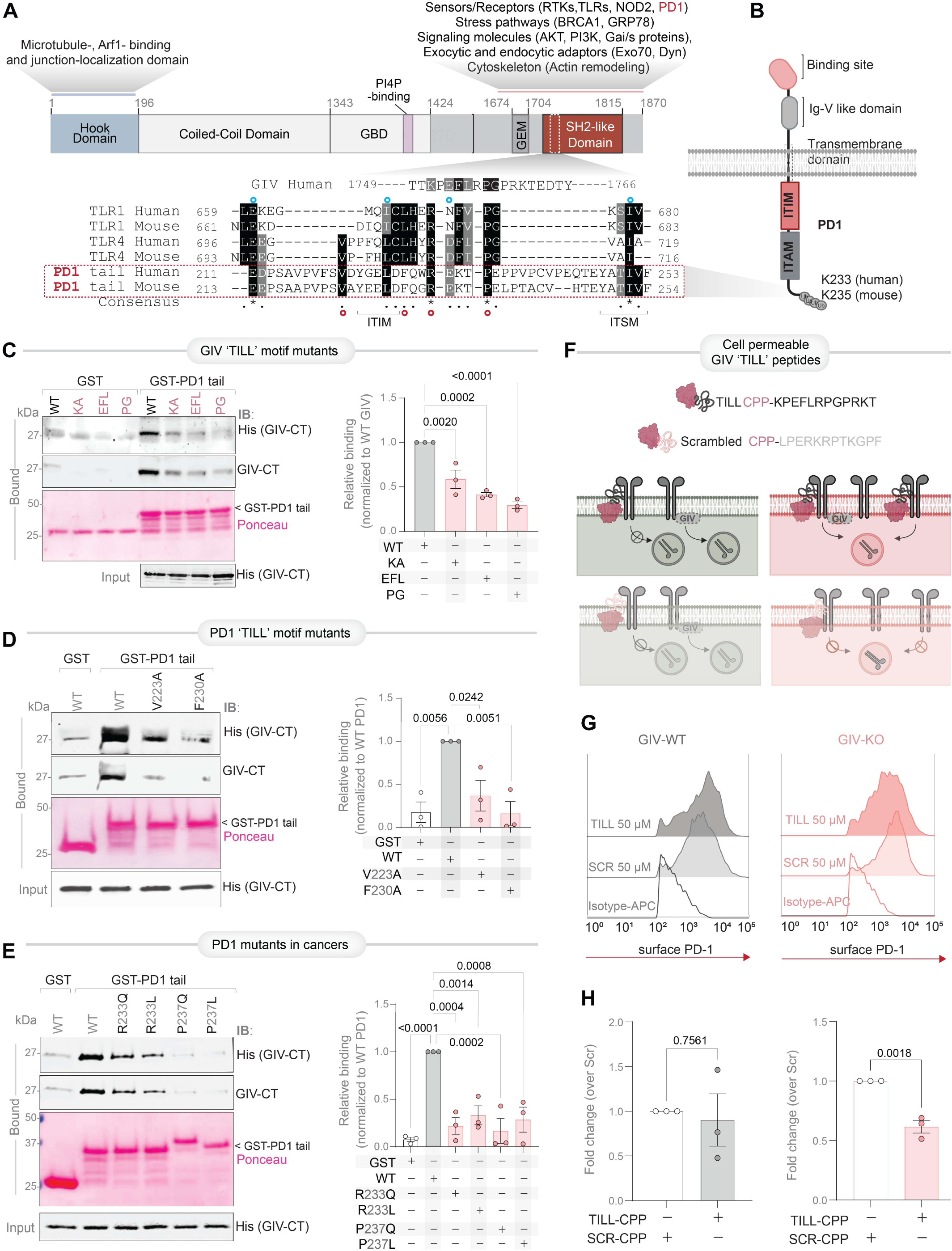
A conserved TILL motif enables GIV–PD1 coupling to control receptor trafficking. **(A, B)** Domain architecture of human GIV and PD1, highlighting the C-terminal region of GIV and the cytoplasmic tail of PD1. Sequence alignment (A, lower) identifies a conserved short linear TIR-like loop (TILL) motif shared between GIV, TIR-containing Toll-like receptors (TLRs), and the PD1 cytoplasmic tail, positioning this motif as a candidate interaction interface. Lys(K233), a hotspot for ubiquitination and receptor stabilization^83,84^ is highlighted. Red (**○**) denotes mutant proteins tested in this study, whereas blue (**○**) indicate mutations that failed expression and were excluded from analysis. ITIM, Immunoreceptor Tyrosine-based Inhibitory Motif; ITSM, Immunoreceptor Tyrosine-based Switch Motif. **(C)** GST pulldown assays using recombinant His–GIV C-terminus (CT) demonstrate direct binding to the GST-PD1 tail but not GST alone. Immunoblots (left) and quantification (right) show the impact of mutation of key residues within the GIV’s TILL motif on PD1 binding. GST proteins are visualized by Ponceau S staining. **(D, E)** GST pulldown assays testing PD1 tail mutants. Immunoblots (left) and quantification (right) show the impact of mutation of key residues within the PD1’s TILL motif, including residues frequently mutated in cancer (COSMIC; see **Figure S7A-B**) on GIV binding. **(F)** Schematic of cell-permeable TILL peptides (TILL-CPP) or scrambled control (SCR-CPP) used to competitively perturb GIV–PD1 coupling and assess effects on PD1 endocytic routing. **(G-H)** Flow cytometry histograms (G) and bar plots (H) compare the abundance of surface PD1 upon treatment with TILL-CPP in GIV-KO and WT cells compared to controls (SCR-CPP-treated baseline). *Statistics:* Data are mean ± SEM (n = 3–5 biological replicates). Statistical significance was assessed by Bonferroni’s multiple comparisons test and unpaired t-test, as appropriate. *p*-values < 0.05 were considered significant.

Biochemical reconstitution confirmed this mechanism of direct coupling. Recombinant His–GIV C-terminus bound the GST–PD1 tail, but not GST alone, and mutation of key residues within GIV’s TILL motif disrupted PD1 binding (**Figure 7C**). Reciprocal mutagenesis of the PD1 tail similarly impaired GIV association, including substitutions that mirror recurrent somatic mutations in human cancers (COSMIC) clustering within a membrane-proximal cytosolic patch encompassing the TILL region (**Figure 7D–E**; **Figure S7A–B**). These data define a bilateral TILL•TILL interface as the molecular determinant of specific GIV•PD1 coupling.

To test the functional *sufficiency* of this mode of coupling, we used cell-permeable synthetic peptides (CPPs) encoding the TILL-motif (TILL-sequence fused to the C-terminus of the Antennapedia-CPP^81^, to acutely perturb this interaction (**Figure 7F**). These peptides, representing the minimal required GIV-TILL segment, were predicted to bind PD1. If the TILL motif is necessary, but not sufficient, the peptide would competitively ‘displace’ endogenous GIV in WT cells and inhibit PD1 endocytosis, functioning as a dominant negative perturbagen. If it is required and sufficient, it would facilitate PD1 endocytosis in GIV-KO cells by ‘rescuing’ the genetic defect. Compared to scrambled (SCR) control peptides, acute TILL-CPP treatment did not change surface PD1 in WT macrophages but reduced surface PD1 abundance in GIV-KO cells (**Figure 7G**). Thus, the TILL motif alone is sufficient to engage PD1 and restore endocytic routing in the absence of full-length GIV. Structure-guided homology modeling further supported these findings, revealing a complementary bilateral TILL•TILL interface and highlighting key residues that destabilize this interaction when mutated in cancer (**Figure 8A**).

**Figure 8.**
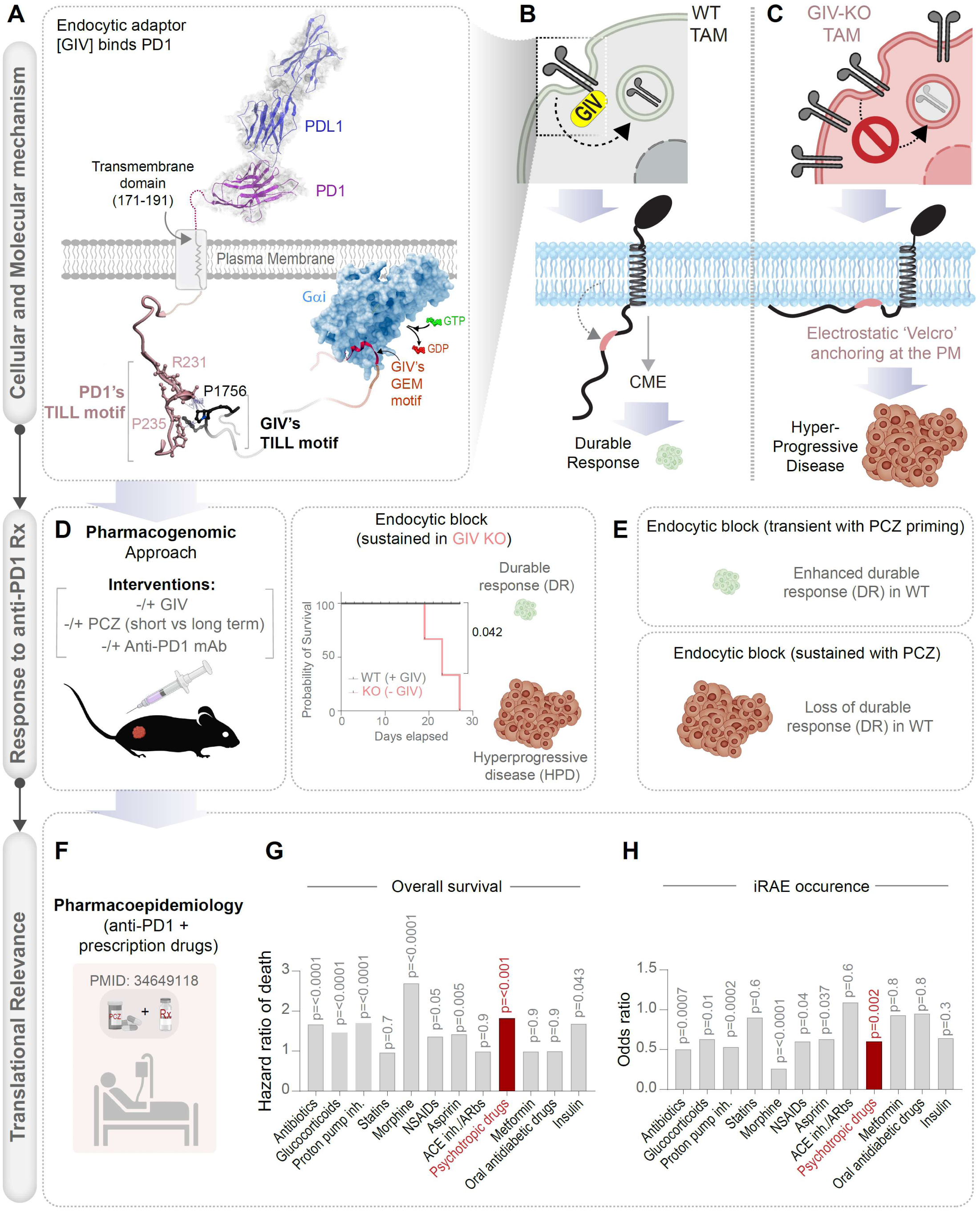
Working model for how GIV-dependent PD1 positioning in TAMs governs immunotherapy durability. **(A)** A homology model of bilateral TILL•TILL binding between the PD1 cytoplasmic tail and the GIV C-terminus (see *Methods*) that is experimentally validated highlighting a conserved PD1•GIV interaction interface. **(B)** In WT TAMs, GIV binds to PD1, couples PD1 to clathrin-mediated endocytosis, limiting surface PD1 accumulation, preserving phagocytic function, and supporting durable responses to PD1 blockade. **(C)** In GIV-deficient TAMs, PD1 is retained at the plasma membrane, suppressing macrophage effector function and driving hyperprogressive disease following PD1 therapy. **(D–E)** Genetic loss of GIV or sustained pharmacologic inhibition of endocytosis phenocopies GIV-KO, whereas transient endocytic modulation enhances PD1 blockade efficacy in GIV-proficient settings. **(F–H)** Reanalysis of pharmacoepidemiologic evidence across multiple cancer types^82^ (F) link concomitant use of endocytosis-inhibiting psychotropic drugs with higher death rates (G) with a concomitant lower incidence of irAEs (Immune-Related Adverse Events; H) in patients receiving anti–PD1/PD-L1 therapy. *Statistics:* Statistical significance of overall survival hazards ratios were assessed using univariable Cox regression and p values were calculated using Fisher’s exact test. *p*-values < 0.05 were considered significant.

These data define a conserved TILL-mediated interaction that couples PD1 to GIV and enables receptor endocytosis and supports a working model for how endocytic control of surface PD1 by GIV represents a central determinant of immunotherapy success (**Figure 8B**). Disruption of this interface, either by genetic deletion, pharmacologic manipulation, mutation or competitive inhibition, traps PD1 at the cell surface. These mechanistic insights link innate immune signaling logic, cancer-associated PD1 mutations, and the durability of immunotherapy responses (**Figure 8C-E**). Pharmacoepidemiologic analyses (**Figure 8F**) mirror our mechanistic findings: concomitant use of such agents during PD1 blockade are associated with worse survival (**Figure 8G**) but reduced immune-related adverse events (**Figure 8H**). Together, our findings provide mechanistic explanations for antibody^40^– and drug-dependent^82^ variability in clinical outcomes.

## DISCUSSION

This study identifies PD1 endocytosis, not ligand blockade alone, as a macrophage-intrinsic immune checkpoint that determines immunotherapy durability. We show that the endocytic adaptor GIV/Girdin binds PD1 through a conserved TIR-like loop (TILL) interaction, enabling clathrin-mediated internalization that limits surface PD1 on TAMs. This routing preserves macrophage phagocytosis and safeguards durable responses to PD1 blockade. Genetic or pharmacologic disruption of this pathway traps PD1 at the plasma membrane, collapses macrophage effector function, accelerates tumor growth, and converts checkpoint therapy into hyperprogressive disease. Thus, macrophage endocytic routing represents a tractable axis for improving the durability of cancer immunotherapy.

### TAM states dictate immunotherapy fate

By integrating a macrophage systems model with single-cell tumor transcriptomes, we define TAM subtypes that stratify durable response versus secondary resistance with accelerated tumor growth at relapse. These states are governed by process-level wiring—innate sensing, endocytic competence, and phagocytic capacity, rather than lineage abundance. GIV serves as a nodal regulator coupling these programs, providing a unifying explanation for TAM heterogeneity. This framework reconciles prior context-dependent observations, including the evolving biology of TREM2⁺ macrophages and their context-specific role in tumors.

### A TIR-like loop embeds PD1 in innate immune logic

We identified a previously unrecognized TIR-like loop (TILL) motif within the PD1 cytoplasmic tail, revealing that PD1 co-opts conserved innate immune interaction logic. PD1 engages GIV through a TILL•TILL interface analogous to TLR–adaptor signaling, reframing PD1 as embedded within innate immune architecture rather than a purely adaptive checkpoint. Our findings suggest that GIV-mediated PD1 endocytosis may represent a physiological brake-release mechanism, allowing macrophages to recalibrate between tolerance and inflammation in response to contextual cues. In cancer, failure of this routing mechanism locks macrophages into a tolerant, non-phagocytic state, undermining antitumor immunity.

Because GIV binds both TLR4 and PD1 via the same motif, these interactions are likely mutually exclusive, positioning GIV as a molecular switch that integrates innate activation with checkpoint resolution. Moreover, spatial proximity of the TILL motif to PD1 K233 (K235 in mice) further suggests crosstalk between GIV binding and receptor stability that is dictated by dynamic ubiquitination at that site^83,84^.

Somatic mutations within the PD1 cytoplasmic tail—particularly those clustering near the TILL region—may be acquired or functionally redeployed in macrophages through tumor material uptake, nanotubes^85^ or cell fusion^86^, generating “tumor-educated” macrophages with altered checkpoint trafficking. Our findings suggest that such mutations could phenocopy GIV loss by stabilizing PD1 at the macrophage surface, thereby promoting immune tolerance and contributing to tumor progression and immunotherapy resistance.

This mechanistic convergence suggests a model in which GIV occupancy integrates innate activation signals with checkpoint receptor fate, coordinating inflammatory tone, receptor turnover, and immune effector function. Disruption of this coordination—genetically or pharmacologically—uncouples innate sensing from checkpoint resolution.

### GIV as an endocytic adaptor for PD1

Our findings establish GIV as a *bona fide* endocytic adaptor for PD1. GIV directly binds dynamin and is known to selectively gate clathrin-mediated endocytosis of defined cargoes—promoting uptake of receptors such as transferrin and E-cadherin while restraining EGFR and β-integrin. Engagement of PD1 by GIV drives PD1 internalization, and notably, the short TIR-like loop (TILL) motif alone is sufficient to confer this endocytic control, independent of GIV’s other signaling modules. This sufficiency was unexpected given GIV’s large, multimodular architecture and its broad roles in cytoskeletal and signaling networks^51–56,87^.

Mechanistically, GIV may facilitate PD1 endocytosis through two non-exclusive mechanisms. First, the GIV-binding region in PD1 cytosolic tail contains a membrane-proximal hydrophobic patch that may serve as an electrostatic–hydrophobic “Velcro,” stabilizing PD1 at the plasma membrane (**Figure 8B-C**), analogous to mechanisms described for PD-L1^88,89^ (**Figure S7C**). Such patches anchor proteins via a two-pronged mechanism: hydrophobic interactions with lipid tails and electrostatic attraction to anionic membrane lipids (e.g., PIP2, phosphatidylserine) ^90^. This dual-site mechanism enables precise control over where and when proteins bind to membranes, thereby affecting signal-coordinated processes like endocytosis^90^. GIV binding likely disrupts this stabilization, licensing PD1 for endocytic uptake. Second, because GIV oligomerizes through its coiled-coil domain^91^, GIV•PD1 engagement may promote receptor clustering, a well-recognized prerequisite for efficient tumor cell internalization, which in turn promotes tumor immunity^40^. Together, these observations integrate biochemical, structural, and functional data into a coherent model in which GIV couples PD1 to the endocytic machinery to control checkpoint receptor fate.

### Endocytic inhibition as a double-edged sword in cancer care

Our findings reveal a critical paradox in cancer care: Endocytic inhibition can both enhance and undermine anti-tumor immunity, depending on the context. Endocytic inhibitors have been widely explored as adjuvants to improve the efficacy of IgG1-based biologics (e.g., cetuximab, trastuzumab, avelumab) by prolonging receptor–antibody engagement and enhancing ADCC^44^. Our data demonstrate that the same strategy is deleterious in the context of PD1 blockade, where sustained inhibition of endocytosis traps PD1 at the macrophage surface and converts immunotherapy into hyperprogressive disease.

This distinction has immediate clinical relevance. Psychotropic and antiemetic drugs that impair dynamin-dependent endocytosis are widely prescribed in oncology, often without consideration of immunotherapy interactions. In light of our mechanistic findings, pharmacoepidemiologic evidence warrants caution: concomitant use of such agents during PD-1 blockade may blunt the development of durable antitumor immunity and worsen survival.

Together, these observations establish endocytic inhibition as a double-edged sword, underscoring the need for temporal precision and pathway awareness when deploying such agents alongside IgG1-based biologics vs immunotherapy.

### Limitations

This study opens several mechanistic avenues that extend beyond the scope of the present work. GIV is a multimodular scaffold, and we focused on its endocytic adaptor function; how its additional domains couple PD1 to tyrosine kinase pathways or heterotrimeric G protein signaling remains to be defined. Notably, GIV potently suppresses the cAMP–PKA axis^51,59,92^, an emerging driver of protumorigenic TAM states^93–95^, raising the possibility that PD1 trafficking intersects with metabolic and transcriptional checkpoints downstream of cAMP signaling. We did not resolve how GIV dynamically partitions between innate immune receptors (e.g., TLR4, NOD2) and PD1, or how crosstalk between these pathways are temporally coordinated during immune activation. The proximity of the GIV-binding site to PD1 K233 (K235 in mice) highlights an unexplored interface between endocytic routing and ubiquitin-mediated control of receptor stability^83,84^. Whether GIV biases PD1 toward degradation versus recycling remains an open question. Because PD1 and PD-L1 are co-expressed in myeloid cells, the possibility of *cis* interactions^96,97^ at the plasma membrane—and GIV’s role in such signaling—remains unexplored. Finally, as GIV is also expressed in T cells^98^, its endocytic functions may extend to PD1 positioning in adaptive immunity.

Despite these gaps, this work reframes PD1 trafficking as a macrophage-encoded determinant of immunotherapy durability, providing mechanistic resolution to long-standing clinical paradoxes and defining actionable principles for improving the safety and durability of cancer immunotherapy. By linking PD-1 routing to GIV, we position endocytic control as a nexus that integrates signaling, metabolism, and receptor fate, highlighting endocytic control as a rich, tractable frontier for reshaping immunotherapy responses.

## Resource Availability

### Lead Contact

Further information and requests for resources and reagents should be directed to and will be fulfilled by the lead contact, Pradipta Ghosh.

### Materials Availability

All materials are available from the lead contact with a completed Materials Transfer Agreement and patented technology agreement following the guidelines of the University of California, San Diego.

### Data and Code Availability

All transcriptomic datasets generated in this work has been deposited at NCBI GEO [<u>GSE314811</u> and <u>GSE314813</u>; see *KRT*; accessible to reviewers (<u>GSE314811</u> code: mhwdmwoevhepvkf, <u>GSE314813</u> code: ojadwccefludrav) and will be made public at the time of publication]. All code used to compute composite score, AUC and generate the plots is publicly available on GitHub at: https://github.com/sinha7290/Mullick_et_al. The repository includes the BoNE framework scripts and the Jupyter notebook used to reproduce key figure panels. The data underlying all the figures and tables are available in the article and its online supplemental materials. Any additional information required for reanalysing the data reported in this work paper is available from the Lead Contact upon request.

## Supporting information

Supplemental details

## Acknowledgments

This work was supported by the National Institutes of Health (NIH) (grant# R01-AI141630, R01-CA238042 and R01-AI55696 to P.G.). M.M. was supported by a UC San Diego Agilent Center of Excellence Postdoctoral Fellowship. G.D.K. received support from the American Association of Immunologists Intersect Fellowship Program for Computational Scientists and Immunologists. Both G.D.K and M.S.A were also supported in part by the American Heart Association (24CDA1268506 and 25PRE1357971, respectively). S.S. was supported by the AAI Intersect Fellowship Program. P.A.T was supported by NIH-NHLBI R01169861. Data in this manuscript were generated at the UC San Diego Institute of Genomic Medicine using an Illumina NovaSeq 6000, funded by NIH SIG grant S10-OD026929. The authors acknowledge instrumentation resources at the UC San Diego Agilent Center of Excellence in Cellular Intelligence. The authors would like to thank the University of California, San Diego – Cellular and Molecular Medicine Electron Microscopy Core (UCSD-CMM-EM Core, RRID:SCR_022039) for TEM sample preparation and microscope access. The content is solely the authors’ responsibility and does not necessarily represent the official views of the AHA or the NIH.

## Author contributions

M.M and P. G conceptualized the project. S.S. and P. G conceptualized and performed all transcriptomic and systems-level analyses. S.S supervised all biostatistical approaches. E.M., S.R., B.B., V.C., M.S.A., and C.R.E., contributed to data curation and formal analysis. V.C. maintained and genotyped *Ccdc88a*^fl/fl^ *LysM*^Cre/-^ mice and C.R.E conducted tumor tissue transcriptomic analyses. M.M., E.M. and B.B. conducted all animal studies, imaging experiments, and quantitative image analyses. C.R.E. and S.R performed site-directed mutagenesis of all constructs used in this work. S.R. and M.S.A conducted all biochemical and molecular interaction studies. P.A.T. procured surgically resected lung tissues, which served as the source material for organoid derivation and benchmarking by C.T. and S.W. M.M. and P.G. prepared the Figures for data visualization. PG wrote the original draft. M.M., E.M., S.R., C.R.E., G.D.K., S.S., and P.G. reviewed and edited the manuscript. P.G. supervised the study and secured funding. All authors approved the final version of the manuscript.

## STAR* METHODS

KEY RESOURCES TABLE.

| REAGENT or RESOURCE | SOURCE | IDENTIFIER |
| --- | --- | --- |
| **Biological Samples and Cell Lines** |  |  |
| RAW 264.7 | ATCC | TIB-71 |
| shControl RAW 264.7 | Swanson et al. 2019<sup>49</sup> |  |
| shGIV RAW 264.7 | Swanson et al. 2019<sup>49</sup> |  |
| HeLa parental | ATCC | ATCC® CCL-2 |
| HeLa GIV KO (CRISPR Cas9) | Lab generated <sup>102,103</sup> | n/a |
| Lewis Lung Carcinoma | ATCC |  |
| THP-1 human monocyte | ATCC | TIB-202 |
| *LysMcre* mice (*B6.129P2-Lyz2<sup>tm1(cre)lfo/j</sup>*) | The Jackson Laboratory | 004781 |
| Girdin flox mice | Asai et al. 2012 <sup>104</sup> | N/A |
| *Ccdc88a<sup>fl/fl</sup>* *LysM<sup>Cre/-</sup>* mice | Swanson et al. 2019<sup>49</sup> |  |
| **Plasmids and constructs** |  |  |
| His GIVCT WT | Swanson et al. 2019<sup>49</sup> | N/A |
| His GIVCT-KA | Swanson et al. 2019<sup>49</sup> | N/A |
| His GIVCT-EFL | Swanson et al. 2019<sup>49</sup> | N/A |
| His GIVCT-PG | Swanson et al. 2019<sup>49</sup> | N/A |
| GST-EMPTY vector |  | N/A |
| PD1-GFP (mouse) | Peled M. et al. 2018<sup>105</sup> | N/A |
| GST-PD1-CT (mouse) | Peled M. et al. 2018<sup>105</sup> | N/A |
| GST-PD1-V223A | *This study* | Amino acid # per mouse REFSEQ: accession NM_008798.3 |
| GST-PD1-F230A | *This study* | Amino acid # per mouse REFSEQ: accession NM_008798.3 |
| GST-PD1-R233Q | *This study* | Amino acid # per mouse REFSEQ: accession NM_008798.3 |
| GST-PD1-R233L | *This study* | Amino acid # per mouse REFSEQ: accession NM_008798.3 |
| GST-PD1-P237Q | *This study* | Amino acid # per mouse REFSEQ: accession NM_008798.3 |
| GST-PD1-P237L | *This study* | Amino acid # per mouse REFSEQ: accession NM_008798.3 |
| pET-28b-GIV-CT-WT (a.a. 1660-1870) | Garcia-Marcos et al 2009<sup>58</sup> | Referred to as His-GIV-CT (aa 1660-1870) |
| pGEX-4T-GIV-CT-WT (a.a. 1623-1870) | Garcia-Marcos et al 2009<sup>58</sup> | Referred to as GST-GIV-CT (aa 1660-1870) |
| pCEFL-GST-GIV-CT (1660-1870) | Bhandari et al 2015<sup>106</sup> | Referred to as pCEFL GST-GIV-CT for mammalian expression (aa 1660-1870) |
| CMV14-p3X FLAG-GIV WT (full length) | Garcia-Marcos et al 2009<sup>58</sup> | N/A |
| **Chemicals and Reagents** |  |  |
| TRIzol™ Reagent | Life Technologies | Cat#15596018 |
| Quick-RNA MiniPrep Kit | Zymo Research | Cat#R2052 |
| qScript™ cDNA SuperMix | QuantaBio | Cat#101414 |
| PowerUp™ SYBR™ Green Master Mix | Applied Biosciences | Cat#A25741 |
| Applied Biosystems StepOnePlus Real Time PCR System Spectral Calibration Kit (96 well) | Thermo Fisher Scientific | Cat#4371435 |
| Direct-zol RNA Miniprep Kit | Zymo Research | Cat#R1051 |
| Zinc Formalin Fixative | Sigma-Aldrich | Cat#Z2902 |
| Phorbol 12-myristate 13-acetate | Millipore Sigma | Cat#P1585 |
| pHrodo Red succinimidyl ester (pHrodo-SE) | Invitrogen | Cat# P36600 |
| CFSE dye | Invitrogen | Cat#C1157 |
| CellTracker Deep Red Dye | Invitrogen | Cat#C34565 |
| CellTracker™ CM-Dil Dye | Invitrogen | Cat#C7001 |
| RPMI 1640 media with L-glutamine | Cytiva | Cat#SH30027.FS |
| RPMI 1640 media, no phenol red | Gibco | Cat#11-835-055 |
| DMEM/High glucose with L-glutamine; without sodium pyruvate | Cytiva | Cat#SH30022.FS |
| Fetal Bovine Serum | Sigma-Aldrich | Cat#F8067-500ML |
| Penicillin Streptomycin 100X Solution | Cytiva | Cat#SV30010 |
| Trypsin-EDTA (0.25%), phenol red | Fisher Scientific | Cat#25-200-114 |
| Dulbecco's Phosphate Buffered Saline (DPBS), 1X, without calcium, magnesium | Cytiva | Cat#16750-076 |
| Matrigel | Corning | Cat#354234 |
| Prochlorperazine dimaleate salt | Selleck Chemicals | Cat#S4631 |
| Mayer's hematoxylin | BioGenex | Cat#HK1005K-GP |
| Two-component DAB brown | BioGenex | Cat#HK542XAKE-GPE |
| Collagenase I | GIBCO | Cat#17100-017 |
| DNase I | Zymo Research | Cat#E1011-A |
| Human TruStain FcX™ | BioLegend | Cat#422302 |
| Poly-L-lysine coated slides | Fisher Scientific | Cat#63410-02 |
| 3% hydrogen peroxide | Fisher Scientific | Cat#S25359 |
| Dyngo4a | Abcam | Cat#AB120689 |
| Duolink® In Situ Detection Reagents Red | Sigma-Aldrich | Cat#DUO92008-30RXN |
| Duolink® In Situ PLA® Probe Anti-Rabbit PLUS, Affinity purified Donkey anti-Rabbit IgG (H+L) | Sigma-Aldrich | Cat#DUO92002-30RXN |
| Duolink® In Situ PLA® Probe Anti-Mouse MINUS, Affinity purified | Sigma-Aldrich | Cat#DUO92004-30RXN |
| Donkey anti-Mouse IgG (H+L) |  |  |
| Fluid Thioglycollate Medium | BD Difco | Cat#225650 |
| 2% paraformaldehyde | Electron Microscopy Services | Cat#15700 |
| 0.2% glutaraldehyde | Electron Microscopy Services | Cat#16200 |
| 0.1% Probumin® BSA | Millipore Sigma | Cat#820452 |
| 2% gelatin | Sigma-Aldrich | Cat# ES-006-B |
| 12 nm Colloidal Gold | Jackson Immuno research lab | Cat#115-205-068 |
| 18 nm Colloidal Gold | Jackson Immuno research lab | Cat#115-215-068 |
| 4% uranyl acetate | Polysciences | Cat#21447-25 |
| 2% methylcellulose | Sigma Aldrich | Cat#9004-67-5 |
| Tyr Phosphatase inhibitor cocktail | Sigma-Aldrich | P5726 |
| Ser/Thr phosphatase inhibitor cocktail | Sigma-Aldrich | P0044 (replaced with P2850 later) |
| Protease inhibitor cocktail | Roche Diagnostics | Cat#5056489001 |
| Glutathione Sepharose beads | Sigma-Aldrich | Cat#17-0756-04 |
| Antibodies |  |  |
| Anti-polyhistidine antibody (monoclonal) | Sigma-Aldrich | Cat#H1029-.2ML |
| 16% Paraformaldehyde | Electron Microscopy Sciences | Cat#15710 |
| ProLong Glass Antifade Mountant | Thermo Fisher Scientific | Cat#P36984 |
| Pierce™ Protein G Agarose | Thermo Fisher Scientific | Cat# 20398 |
| Limonene Mounting Medium | Abcam | Cat#ab104141 |
| 2.5% Normal Goat Serum | Vector Laboratories | Cat#S-1012-50 |
| 2.5% Normal Horse Serum | Vector Laboratories | Cat#S-2012-50 |
| IPTG | Zymo Research | Cat# I1001-25 |
| **Antibodies** |  |  |
| InVivoMab anti-mouse PD1 (CD279, monoclonal) (Clone: RMP1-14) | Bio X Cell | Cat#BE0146 |
| PD1 Antibody (polyclonal) | Proteintech | Cat#18106-1-AP |
| PD1 Antibody (monoclonal) | Invitrogen | Cat#MA5-15780 |
| Recombinant Mouse B7-H1 (PD-L1, CD274)-Fc Chimera (carrier-free) | BioLegend | Cat#758208 |
| Anti-NCAM1 (CD56) antibody (monoclonal) | Abcam | Cat#ab220360 |
| Anti-CD68 antibody (monoclonal) | Abcam | Cat#ab201340 |
| CD3e antibody (monoclonal) | eBioscience, Invitrogen | Cat#14-0031-82 |
| Anti-CD163 antibody (monoclonal) | Abcam | Cat#ab182422 |
| Anti-Girdin, coiled-coil region antibody (polyclonal) | Millipore Sigma | Cat#ABT80 |
| CK7 antibody (monoclonal) (OVT12/30) | Invitrogen | Cat#MA1-06316 |
| Impress™ HRP Anti-Rabbit IgG Polymer Detection Kit, Peroxidase | Vector Laboratories | Cat#MP-7401-50 |
| Impress™ HRP Anti-Mouse IgG Polymer Detection Kit, Peroxidase | Vector Laboratories | Cat#MP-7402-50 |
| Goat anti Rabbit IgG (H+L) Secondary Antibody, Alexa Fluor 647 | Invitrogen | Cat#A-21245 |
| Goat anti-Syrian Hamster IgG (H+L) Cross-Absorbed Secondary Antibody, Alexa Fluor 488 | Invitrogen | Cat#A-21110 |
| F(ab')<sub>2</sub>-Goat anti-Rabbit IgG (H+L) Cross-Absorbed Secondary Antibody, Alexa Fluor 594 | Invitrogen | Cat#A-11072 |
| Zombie UV™ Fixable Viability Kit | BioLegend | Cat#423108 |
| PerCP anti-mouse/human CD11b antibody (monoclonal) | BioLegend | Cat#101230 |
| FITC anti-mouse F4/80 antibody (monoclonal) | BioLegend | Cat#123108 |
| APC anti-mouse CD279 (PD1) antibody (monoclonal) | BioLegend | Cat#135210 |
| Coralite Plus 488-conjugated CD68 antibody (polyclonal) | Proteintech | Cat#CL488-2574 |
| PE/Cyanine7 anti-mouse CD163 antibody (monoclonal) | BioLegend | Cat#155320 |
| APC/Cyanine7 anti-mouse CD49b (pan-NK cells) antibody (monoclonal) | BioLegend | Cat#108920 |
| FITC anti-mouse CD3 antibody (monoclonal) | BioLegend | Cat#100204 |
| PerCP anti-mouse CD4 antibody (monoclonal) | BioLegend | Cat#100432 |
| APC anti-mouse CD8a antibody (monoclonal) | BioLegend | Cat#100712 |
| PerCP/Cyanine5.5 anti-mouse CD45 antibody (monoclonal) | BioLegend | Cat#103132 |
| FITC anti-mouse Ly-6C antibody | BioLegend | Cat#128006 |
| APC anti-mouse Ly-6G antibody (monoclonal) | BioLegend | Cat#127614 |
| **Deposited data** |  |  |
| Transcriptomic dataset, syngeneic whole tumors | *This paper* | NCBI GEO: GSE314811 |
| Transcriptomic dataset, F4/80 magnetic sorted Tumor Associated Macrophages (TAMs) | *This paper* | NCBI GEO: GSE314813 |
| **Software** |  |  |
| ImageJ | National Institute of Health | https://imagej.net/Welcome |
| QuPath v0.6.0 | Open Source | https://qupath.github.io/ |
| Gen5 | Agilent | https://www.agilent.com/en/product/cell-analysis/cell-imaging-microscopy/cell-imaging-microscopy-software/biotek-gen5-software-for-imaging-microscopy-1623226 |
| Prism 9 | GraphPad | https://www.graphpad.com/scientific-software/prism/ |
| Molsoft | Molsoft, LLC | https://www.molsoft.com/index.html |
| LAS-X | Leica | www.leica-microsystems.com/products/microscope-software/p/leica-las-x-ls |
| SnapGene | GSL Biotech LLC | https://www.snapgene.com/ |
| Illustrator | Adobe | https://www.adobe.com/products/illustrator.html |
| ImageStudio Lite | LI-COR | https://www.licor.com/bio/image-studio-lite/ |
| ClueGO | Cytoscape | https://academic.oup.com/bioinformatics/article/25/8/1091/324247 |
| Python3 | Python | https://www.python.org |
| matplotlib | Matplotlib | https://matplotlib.org/ |
| Seaborn 0.12 | Seaborn | https://seaborn.pydata.org |
| Photoshop | Adobe | https://www.adobe.com/products/photoshop.html |

## DETAILED EXPERIMENTAL METHODS

### Cell culture

Thioglycolate-elicited murine peritoneal macrophages (TGPMs) were isolated from 8-12-wk-old C57BL/6 mice by peritoneal lavage with 10 ml of ice-cold RPMI (Roswell Park Memorial Institute Medium) per mouse, 4 days after intraperitoneal injection of 2.5 ml of aged, sterile 3% thioglycolate broth (BD Difco, NJ, USA) and cultured as described previously^107^. Cells were passed through a 70 µm filter to remove tissue debris, counted, centrifuged, and resuspended in RPMI-1640 containing 10 % FBS and 1% penicillin/streptomycin. Cultures were maintained at 37°C in a humidified 5% CO_2_ incubator. Cells were plated at the required density, and the medium was replaced after 4 hours to remove non adherent cells. Cells were allowed to equilibrate overnight before various stimulations and/or treatments, as indicated in the figure legends.

Lewis lung carcinoma (LL/2), HeLa, THP-1 monocytes, and RAW264.7 cells were obtained from American Type Culture Collection (ATCC). RAW264.7 and THP-1 cells were maintained in RPMI media, while LL/2 and HeLa cells were maintained in DMEM (Dulbecco’s Modified Eagle Medium); all supplemented with 10% FBS and 1% penicillin/Streptomycin. All macrophages were grown in serum that was heat inactivated (at 56°C for 30 minutes).

### Collection of human lung specimens for organoid isolation

Adult lung organoids—from both lung adenocarcinomas and adjacent normal tissues--were generated, as outlined previously^108^, from fresh surgical resections, prospectively collected by a cardiothoracic surgeon. Prior to collection of the lung specimens, each tissue was sent to a gross anatomy room where a pathologist cataloged the area of focus, and the extra specimens were routed to the research laboratory in Tissue-2-Go (HUMANOID^TM^ Cat# HUM202403) for cell isolation. Deidentified lung tissues obtained during surgical resection, which were deemed excess by clinical pathologists, were collected using an approved human research protocol (IRB#101590; PI: Patricia Thistlewaite). Isolation and biobanking of organoids from this lung tissue were carried out using an approved human research protocol (IRB# 190105; PI: Ghosh) that covers human subject research at the UC San Diego HUMANOID Centre of Research Excellence (CoRE). For all de-identified human subjects’ information, including age, gender, and previous history of the disease, was collected from the chart following the rules of HIPAA and described in **Table 1**.

### Isolation and culture of human whole lung-derived organoids

A previously published protocol^109,110^ was modified^108^ to isolate lung organoids from three human subjects. Briefly, normal human lung specimens were washed with PBS containing 4× penicillin-streptomycin and minced using surgical scissors. The resulting tissue fragments were resuspended in 2 ml of wash buffer (Advanced DMEM/F-12, 10 mM HEPES, 1× Glutamax, 1× penicillin-streptomycin) containing 2 mg/ml Collagenase Type I (Thermo Fisher, MA, USA) and incubated at 37°C for approximately 1 hr. During incubation, the tissue was mechanically dissociated every 10 minutes using a 1 ml micropipette and monitored under a light microscope. Once 80–100% of the cells were released from connective tissue, digestion was halted by adding 10 ml of wash buffer supplemented with 2% fetal bovine serum. The suspension was then passed through a 100 µm cell strainer and centrifuged at 200 rcf.

Residual erythrocytes were lysed using 2 ml of red blood cell lysis buffer (Invitrogen, MA, USA) at room temperature for 5 min, followed by the addition of 10 ml of wash buffer and another centrifugation step at 200 rcf. The resulting cell pellets were resuspended in cold Matrigel (Corning,NY, USA) and seeded in 25 µl droplets on a 12-well tissue culture plate. The plate was inverted and incubated at 37°C for 10 min to allow complete polymerization of the Matrigel before the addition of 1 ml Lungbrew (HUMANOID^TM^ cat# HUM2020-250) per well.

Organoids were maintained in a humidified incubator at 37°C/5% CO_2_, with a complete media change performed every 3 days. After 7 and 10 days, when the desired confluency was reached, organoids were harvested using PBS with 0.5 mM EDTA and centrifuged at 200 rcf for 5 minutes. They were then dissociated in 1 ml of TrypLE Select (Gibco, USA) per well at 37°C for 4–5 minutes, followed by mechanical shearing. Wash buffer was added at a 1:5 ratio (TrypLE to wash buffer), and the cell suspension was centrifuged, resuspended in Matrigel, and re-seeded at a 1:5 split ratio.

Lung organoids were biobanked, and passages 3–10 were used for experiments. Subculturing was performed every 7–10 days.

### Mice

Girdin floxed mice were a generous gift from Dr. Masahide Takahashi (Nagoya University, Japan) and were developed as previously described^9^. LysMcre mice (B6.129P2-Lyz2tm1(cre)lfo/j) were obtained from the Jackson Laboratory. Girdin floxed x LysMcre mice were generated previously in our laboratory as described^49^ and were maintained as homozygous floxed (fl/fl) with heterozygous expression of LysM. Primers used for genotyping are listed in *Key Resource Table (KRT)*.

All mouse studies were approved by the University of California, San Diego Institutional Animal Care and Use Committee (IACUC). Mice were housed in the UC San Diego animal facility under a 12-hour light/dark cycle (30–70% humidity, room temperature maintained at 68–75 °F) with ad libitum access to standard chow and water.

### Sex as a biological variable

Sex was not considered as a biological variable in this study. Both male and female mice were used.

### Establishment of syngeneic ectopic tumor model in mice and drug treatment

Mice were injected subcutaneously on the right hindlimb flank with 0.5 x 10^6^ Lewis lung cancer cells suspended in 0.1 ml of sterile phosphate buffered saline (PBS) supplemented with 20% Matrigel.

Drug treatments started when tumors reached approximately 50 mm^3^ and mice were sacrificed at a maximum tumor volume of 2000 mm³, which was considered the study endpoint. Mice were weighed prior to the start of drug treatment, and tumor dimensions were measured every alternate day using a digital caliper (Vernier, Silverline Digital) for a total of three weeks. Tumor volume in each mouse was calculated using the formula: **Tumor Volume** = (*length x width*^2^)/2

### Drug Treatment Regimens

- *Anti-PD1 mAb monotherapy*: Upon reaching palpable tumor volume, mice received anti-PD1 monoclonal antibody (InVivo mAb, BioXcell) at a concentration of 1 mg/ml in sterile PBS. A volume of 0.1 ml was administered intraperitoneally per mouse, every 3 days, for a total of three doses, as described previously^111^.
- *PCZ monotherapy*: The dynamin inhibitor prochlorperazine dimaleate (PCZ; Sigma-Aldrich, St Louis, MO)^44^ was weighed and dissolved in 1 ml of Dimethyl Sulfoxide (DMSO) (Sigma-Aldrich). A working stock solution of PCZ (1 mg/ml) was prepared using PBS in the biological safety cabinet prior to injection. Mice in the PCZ and PCZ+ anti-PD1 mAb groups received PCZ at a final concentration of 0.8 mg/kg.
- *PCZ-primed PD1 therapy*: Upon palpable tumor volumes were achieved, animals were injected intraperitoneally with PCZ (0.8–1 mg/kg). Twenty-four hours later, anti-PD1 mAb treatment was initiated and administered every three days for a total of three doses over three weeks.

All treatments were administered via intraperitoneal injection in a total volume not exceeding 120 µl. Tumor growth inhibition was assessed by comparing tumor volumes between groups using two-way ANOVA in GraphPad Prism (Dotmatics Inc., Boston, MA).

### Tissue histology

Histological processing of excised tumors was performed at the Histology Core Facility of the La Jolla Institute for Immunology, La Jolla, California. Tissues were initially fixed in zinc formalin for 24 hours, followed by transfer to 70% ethanol and storage at room temperature until sectioning. Samples were then sectioned for hematoxylin and eosin (H&E) staining, and adjacent unstained sections were processed for immunohistochemical (IHC) staining using markers of interest. Stained sections were scanned and imaged using a Leica light microscope (Leica Microsystems, Wetzlar, Germany), and the resulting images were analyzed using ImageJ/Fiji software (NIH, Bethesda, MD)^112^.

### Immunohistochemical and immunofluorescent staining of tumor tissue sections

Harvested tumor specimens were fixed in zinc paraformaldehyde, followed by transfer to 70% ethanol for preparation of formalin-fixed, paraffin-embedded (FFPE) tissue blocks. Sections of 4 μm thickness were cut and mounted onto poly-L-lysine–coated glass slides, followed by deparaffinization and rehydration. Heat-induced epitope retrieval was performed using sodium citrate buffer (pH 6.0) or Tris-EDTA buffer (pH 9.0), depending on the target immune marker, in a pressure cooker.

Sections were then treated with 3% hydrogen peroxide for 10 minutes to quench endogenous peroxidase activity and incubated overnight at 4°C with primary antibodies in a humidified chamber^113^. The following primary antibodies were used for immunohistochemistry (IHC):

- PD1, rabbit polyclonal (1:100 dilution)
- CD3, Armenian Hamster monoclonal (anti-mouse, 1:50 dilution)
- CD163, rabbit polyclonal (1:150 dilution)
- CD56, rabbit polyclonal (1:50 dilution)

Signal detection was carried out using a streptavidin–biotin complex and 3,3′-diaminobenzidine (DAB) as the chromogen, followed by hematoxylin counterstaining. Images were acquired using a Leica light microscope (Leica Microsystems, Wetzlar, Germany) and analyzed using Fiji/ImageJ software (NIH, Bethesda, USA).

### Immunofluorescence Staining

For immunofluorescence double staining, additional FFPE sections were processed similarly with antigen retrieval. The following primary antibodies were used:

- CD3, Armenian Hamster monoclonal (anti-mouse, 1:50 dilution)
- CD68, mouse monoclonal (1:150 dilution)
- CD163, rabbit polyclonal (1:150 dilution)
- CD56, rabbit polyclonal (1:50 dilution)

Secondary antibodies included donkey anti-Goat IgG (H+L) conjugated to Alexa Fluor 488 and Alexa Fluor 594 (1:500 dilution), and nuclei were counterstained with DAPI (1:1000 dilution). Fluorescent images were acquired on a Stellaris 5 Confocal Microscope (Leica Microsystems) and analyzed using QuPath (v0.1.2), with further processing in Fiji/ImageJ.

### Sample collection and assessment of immune cell populations by flow cytometry

Tumors were harvested and weighed; a small portion was fixed in formalin for histological analysis. The remaining tissue was finely minced, homogenized, and digested at 37 °C for 25 minutes in digestion media consisting of RPMI (serum-free) supplemented with 28 U/ml Collagenase I and 10 mg/ml DNase I, as previously^114^.

The resulting cell suspension was passed through a 100 μm strainer, and the strainer was washed with 5–10 ml of complete RPMI medium (containing 10% heat-inactivated FBS, 100 U/ml penicillin, 100 μg/ml streptomycin, and non-essential amino acids). The suspension was then filtered through a 70 μm strainer to obtain a final single-cell preparation. These steps were done as described previously^115^.

Mouse Fc receptors were blocked using TruStain FcX (BioLegend, San Diego, CA) at a 1:500 dilution in PBS (GIBCO) with 1% BSA for 15 minutes. Surface staining was performed using combinations of fluorescently labeled monoclonal antibodies (listed in the Key Resources Table), diluted in PBS containing 1% BSA, and incubated for 30 minutes to 1 hour on ice in the dark. Cells were then washed once with PBS containing 1% BSA to remove unbound antibodies and resuspended in 1 ml of the same buffer for flow cytometric analysis.

Flow cytometry was performed on a NovoCyte Quanteon (Agilent Technologies, Santa Clara, CA), and data were analyzed using NovoExpress (Agilent) and FlowJo software. Gating for positive and negative populations was established using Fluorescence Minus One (FMO) controls, and PD1 gating was guided by appropriate isotype controls. Gating strategies for immune cell populations were based on established protocols and are outlined below:

- **Tumor-Associated Macrophages (TAMs):** Zombie-negative, CD11b⁺, F4/80⁺, PD1⁺
- **Inflammatory TAM Subset:** Zombie-negative, CD11b⁺, CD68⁺, PD1⁺
- **Tolerant TAM Subset:** Zombie-negative, CD11b⁺, CD163⁺, PD1⁺
- **Natural Killer (NK) Cells:** Zombie-negative, CD49b⁺, CD3⁻, CD56⁺
- **T Cells:** Zombie-negative, CD3⁺, CD4⁺, CD8⁺
- **Neutrophils:** Zombie-negative, CD45⁺, Ly6C^low^⁺, Ly6G^high^⁺

### Immunohistochemical and immunofluorescent staining of tumor tissue sections

Tumor specimens were fixed in zinc paraformaldehyde, followed by fixation in 70% ethanol to prepare formaldehyde-fixed paraffin-embedded (FFPE) tissue blocks. Sections (4 μm thick) were cut and mounted onto poly-L-lysine–coated glass slides, then deparaffinized and rehydrated. Heat-induced epitope retrieval was performed in a pressure cooker using sodium citrate buffer (pH 6.0) or Tris-EDTA buffer (pH 9.0), depending on the immune cell marker being stained. Endogenous peroxidase activity was quenched with 3% hydrogen peroxide for 10 minutes. Slides were then incubated overnight at 4°C in a humidified chamber with primary antibodies, including rabbit polyclonal anti-PD1 (1:100) (used only for immunohistochemical staining), mouse monoclonal anti-CD68 (1:150), Armenian Hamster monoclonal anti-CD3 (1:50), rabbit polyclonal anti-CD163 (1:150), and rabbit polyclonal anti-CD56 (1:50) (used for both immunohistochemical and immunofluorescent staining).

Immunohistochemical staining was visualized using a streptavidin–biotin detection system with 3,3′-diaminobenzidine (DAB) as the chromogen and hematoxylin as the counterstain. Images were captured using a Leica light microscope and quantified using Fiji/ImageJ software (NIH, Bethesda, MD, USA).

For immunofluorescence, double staining was performed on unstained FFPE tissue sections following antigen retrieval. Post-primary antibody incubation corresponding secondary antibodies—donkey anti-goat IgG (H+L) conjugated to Alexa Fluor 488 or Alexa Fluor 594 (1:500)—were used along with DAPI nuclear counterstain (1:1000). Fluorescent images were acquired on a Stellaris 5 confocal microscope (Leica Microsystems), processed using QuPath (version 0.1.2), and further analyzed with Fiji/ImageJ software (NIH, Bethesda, MD, USA).

### *In vivo* phagocytosis assay

Eight– to ten-week-old mice were injected intraperitoneally with 3 mL of 3% (w/v) thioglycolate medium. Four days later, 5 × 10⁶ CFSE-labeled LL/2 tumor cells, resuspended in 200 μL of sterile PBS, were administered intraperitoneally. Peritoneal cells were harvested by lavage with ice-cold RPMI medium at 4-, 6-, 12-, and 24-hours post-injection.

Phagocytosis of tumor cells was quantified by flow cytometry^116^. The phagocytic percentage was calculated by gating for CFSE⁺ LL/2 tumor cells within the F4/80⁺ macrophage population, as done previously^117^.

Additionally, harvested cells were fixed in 4% paraformaldehyde and visualized by confocal microscopy using the Stellaris 5 Confocal Microscope (Leica Microsystems). The phagocytic index was determined by counting the number of CFSE⁺ tumor cells internalized per 100 macrophages.

### *In vitro* treatment with dynamin inhibitor

For signaling and fluorescence-based analyses, thioglycolate-elicited peritoneal macrophages (TGPMs) and PMA-differentiated THP1 cells were plated at a density of 1 × 10⁶ cells per well on coverslips in 12-well plates. Cells were treated with either 30 μM DynGo4a (a dynamin inhibitor; Abcam, Cambridge, UK) or DMSO (vehicle control) for 30 minutes. Following treatment, cells were washed with ice-cold PBS and stimulated with recombinant mouse B7-H1 (soluble PD-L1/CD274; BioLegend, San Diego, CA) at a concentration of 10 ng/mL for 5 and 10 minutes. Treated cells were then processed for downstream *in vitro* assays.

### *In vitro* phagocytosis assay

There are three key steps:

- *Labeling of Cancer Cells with pHrodo-SE:* As described by Nam *et al.,*2019 LL/2 cancer cells were washed once with sterile PBS to remove serum residues and resuspended at a concentration of 1 × 10⁷ cells in 50 mL PBS. pHrodo-SE dye was added to a final concentration of 120 ng/ml^118^, and the suspension was incubated on a rotator (15 rpm) at room temperature for 30 minutes. After labeling, cells were washed twice with sterile PBS to eliminate unbound dye and resuspended in serum-free, phenol red-free RPMI medium.
- *Quantitative assessment of phagocytosis*: Labeled LL/2 tumor cells (targets) were co-cultured with thioglycollate-elicited peritoneal macrophages (TGPMs, effectors) at a 1:1 target-to-effector ratio. Fluorescence from internalized pHrodo-labeled targets—indicative of phagocytosis in the acidic lysosomal environment—was measured using a fluorescence plate reader (Spark 20M, Tecan) at excitation/emission wavelengths of 505/525 nm as shown previously ^119^.
- *Live imaging of phagocytosis:* For microscopy-based live imaging, 1.8 × 10⁴ PMA-differentiated THP1 macrophages were stained with CM-DiI dye (Invitrogen) and seeded onto fibronectin-coated (10 µg/mL, 45 min at 37 °C) 96-well black clear-bottom microplates and incubated overnight. The following day, patient-derived lung organoid cells were fluorescently labeled with CFSE dye and added at a seeding density of 0.3 × 10⁴ cells per well. Phagocytosis was monitored by time-lapse imaging every 30 minutes for 6 hours using a BioTek Cytation 10 BioSpa and Confocal Imaging Reader (Agilent Technologies).

### Organoid-macrophage co-culture studies

PMA-differentiated THP1 macrophages were fluorescently labeled using CFSE dye. Lung organoids were generated by embedding mixed cells in Matrigel at a seeding density of 2.5 × 10⁴ cells per well and cultured for 7 days. To assess interactions and phagocytic activity, CFSE-labeled THP1 macrophages were added to the mature lung tumor organoids at a 1:1 ratio (2.5 × 10⁴ cells per well) in 8-well chamber slides and co-cultured for 4 hours.

Following co-culture, cells were fixed with 4% paraformaldehyde (PFA) for 30 minutes at room temperature, quenched with 30 mM glycine for 5 minutes, and blocked with PBS containing 1% BSA for 2 hours. Organoids were stained with an anti-CK7 primary antibody (1:200 dilution) and incubated overnight at 4 °C. After washing with PBS, samples were incubated with Alexa Fluor 594-conjugated secondary antibody (1:500 dilution) and counterstained with DAPI (1:1000 dilution).

Coverslips were mounted using ProLong Glass antifade reagent. Confocal images were acquired using a Stellaris 5 Confocal Microscope (Leica Microsystems). Z-stacks were captured with 1 μm intervals between slices across the relevant channels. For each condition, three representative fields were randomly selected for imaging. Maximum intensity projection images were generated by overlaying z-slices, and all image processing was performed using FIJI (ImageJ) software.

### Duolink™ proximity ligation assays (PLA)

In situ interactions between endogenous GIV and PD1 were detected using the Duolink™ Proximity Ligation Assay kit (Olink Biosciences), following the manufacturer’s protocol with previously optimized modifications^53^. Briefly, PMA-differentiated THP1 cells were cultured and fixed with 4% paraformaldehyde, then permeabilized with 0.1% Tween-20 on ice for 30 minutes. After blocking with Duolink blocking solution for 1 hour at room temperature, cells were incubated with primary antibodies overnight at 4 °C.

Following two washes with 5% BSA in PBS, cells were incubated with PLA probes for 1 hour at 37 °C. Ligation and amplification steps were performed to generate PLA signals.

Confocal images were acquired using a Stellaris 5 Confocal Microscope (Leica Microsystems). For high-throughput quantification of interactions across experimental conditions, images were also analyzed using QuPath (version 0.1.2).

### Confocal immunofluorescence

Cells were plated on coverslips and incubated overnight to reach approximately 80% confluency. Following treatment, cells were fixed in 4% paraformaldehyde in PBS for 30 minutes at room temperature, quenched with 0.1 M glycine for 5 minutes, and then blocked and permeabilized with PBS containing 1% BSA for 30 minutes at room temperature.

Primary antibodies were diluted in blocking buffer and incubated overnight at 4 °C. The following dilutions were used: anti-GIV (1:500) and anti-PD1 (1:500). After primary antibody incubation, cells were washed three times (5 minutes each) with PBS containing 1% BSA and 0.1% Triton X-100.

Fluorescently labeled secondary antibodies, Alexa Fluor 488 and Alexa Fluor 594, were used at 1:500 dilution, and nuclei were counterstained with DAPI at 1:1000 dilution. Coverslips were mounted using ProLong™ Glass Antifade Reagent (Thermo Fisher Scientific).

Images were acquired using a Stellaris 5 Confocal Microscope (Leica Microsystems).

### Immunogold electron microscopy (IEM)

Murine TGPMs were plated in 10 cm dishes at a concentration of 10^7^ cells per dish. The following day, TGPMs were treated with 10 µM dynamin inhibitor, DynGo4a (Abcam, MA, USA) for 30min. Post treatment, cells were fixed with 2% paraformaldehyde and 0.2% glutaraldehyde (15700 and #16200 respectively, Electron Microscopy Services, Hatfield, PA, USA) for one hour at 4°C. The fixative was replaced with ice-cold 4% paraformaldehyde with 0.1% Probumin® BSA in PBS. Cells were gently scraped using a soft Teflon scraper (07-200-366, Thermo Fisher), collected, and processed for sectioning.

*Epoxy resin embedding*: Cells were fixed with 2.5% glutaraldehyde in 0.1M sodium cacodylate buffer pH7.4 for 5 minutes at room temperature and for 2 hours on ice, then pelleted in 10% gelatin. After washing with 0.1 M cacodylate buffer, the pellets are postfixed in 1% OsO4 in 0.1 M cacodylate buffer for 1 hr on ice. The cells were stained all at once with 2% uranyl acetate for 1 hr on ice, following which they were dehydrated in graded series of ethanol (50-100%) while remaining on ice. The cells were then subjected to 1 wash with 100% ethanol and 2 washes with acetone (10 min each) and embedded with Durcupan. Sections were cut at 60 nm on a Leica UCT ultramicrotome and picked up on 300 mesh copper grids. Grids were then incubated for 1 hour at room temperature with primary antibodies diluted in 1% BSA in PBS: anti-PD1 mouse monoclonal antibody (1:10; Invitrogen, Cat. no. MA5-15780) and anti-GIV rabbit polyclonal antibody (1:10; Millipore, Cat. no. ABT80). For immunogold labeling, grids were incubated for 1 hour at room temperature with secondary antibodies diluted 1:1 in 1% BSA in PBS: Goat Anti-Mouse IgG conjugated to 12 nm colloidal gold (Jackson ImmunoResearch, Cat. no. 115-205-068) and Goat Anti-Rabbit IgG conjugated to 18 nm colloidal gold (Jackson ImmunoResearch, Cat. no. 115-215-068). Sections were post-stained with 2% uranyl acetate for 5 minutes and Sato’s lead stain for 1 minute. Images were acquired using a JEM-1400Plus transmission electron microscope (JEOL USA Inc., Peabody, MA), operated at 80KeV and equipped with a bottom-mounted 4kx4k camera Gatan One View.

### Constructs and cloning

Mouse full-length PD1-GFP and GST-PD1 (accession <u>NM_008798.3</u>) cytoplasmic domain constructs were generously provided by Dr. Adam Mor (Columbia University Medical Center). All the PD1 cytoplasmic tail (PD1 CT) mutants used in this work were created using this construct as template^105^. Mutant design was guided either by structure and/or alignment of sequences or inspired by somatic mutations in human cancers listed on the Catalog for Somatic Mutations in Cancer (COSMIC database: https://cancer.sanger.ac.uk).

PD1 CT mutants were created using the Quik-change® Site-Directed Mutagenesis Kit (Stratagene,CA, USA) following the manufacturer’s protocol. His-tagged GIV-CT and GIV-CT TIR domain mutants used in this study were previously developed and validated in our laboratory^49^.

### Immunoblotting

Cell lysates were prepared by resuspending cells in HEPES lysis buffer [20 mM HEPES (pH 7.2), 5 mM magnesium acetate, 125 mM potassium acetate, 0.4% Triton X-100, 1 mM DTT], supplemented with 500 µM sodium orthovanadate, phosphatase inhibitors (Sigma), and protease inhibitors (Roche, California, USA). Lysates were passed through a 28G needle on ice, then cleared by centrifugation at 10,000 × g for 10 minutes.

For immunoblotting, proteins were separated by SDS-PAGE and transferred to PVDF membranes. Membranes were blocked with 5% nonfat milk in PBS, incubated with primary antibodies, and subsequently probed with fluorescent secondary antibodies. Detection and quantification were performed using the LI-COR Odyssey imaging system with dual-color infrared imaging.

All images were processed using Fiji/ImageJ (NIH, Bethesda, USA) and assembled for presentation using Adobe Photoshop and Illustrator (Adobe, San Jose, CA, USA).

### Co-immunoprecipitation (Co-IP)

Cells were lysed in cell lysis buffer [20 mM HEPES (pH 7.2), 5 mM magnesium acetate, 125 mM potassium acetate, 0.4% Triton X-100, 1 mM DTT, 0.5 mM sodium orthovanadate, protein tyrosine phosphatase (PTP) inhibitor cocktail, serine/threonine phosphatase inhibitor cocktail, and protease inhibitor cocktail]. Lysates were sheared by passage through a 28G needle and cleared by centrifugation at 10,000 × g for 10 minutes. Cleared supernatants were incubated 4 h at 4 °C with either mouse anti-PD1 antibody or mouse IgG (control). Protein G agarose beads were then added and incubated for 1 hour at 4 °C to capture immune complexes. Beads were washed with the same phosphate wash buffer described above and eluted by boiling in Laemmli buffer.

### Protein expression and purification

His-tagged and GST-tagged recombinant proteins were expressed in *E. coli* strain BL21 and purified as previously described^120^. Bacterial cultures were induced with 1 mM isopropylβ-D-1-thio-galactopyranoside (IPTG) either overnight at 25°C. Bacterial culture (1l) were pelleted after induction and resuspended in 20 ml GST-lysis buffer (25 mM Tris·HCl, pH 7.5, 20 mM NaCl, 1 mM Ethylenediaminetetraacetic acid (EDTA), 20% [v/v] glycerol, 1% [v/v] Triton X-100, 2 × protease inhibitor mixture [Complete EDTA-free; Roche Diagnostics]) or in 20 ml His-lysis buffer (50 mM NaH_2_PO_4_ [pH 7.4], 300 mM NaCl, 10 mM imidazole, 1% [vol/vol] Triton X-100, 2 × protease inhibitor mixture [Complete EDTA-free; Roche Diagnostics]) for GST or His-fused proteins, respectively. Bacterial lysates were sonicated (three cycles, with pulses lasting 30 s/cycle, and with 2 min interval between cycles to prevent heating), centrifuged at 12,000×*g* for 20 min at 4°C. Supernatant (solubilized proteins) were affinity purified on glutathione-Sepharose 4B beads (GE Healthcare, IL, Chicago) or HisPur Cobalt Resin (Pierce), dialyzed overnight against PBS, and stored at −80°C.

### GST pulldown assays

Recombinant GST-tagged proteins—including GST alone (control), GST-PD1 wild-type, and GST-PD1 mutants—were expressed in *E. coli* strain BL21 (DE3) and purified as previously described^58^. Bacterial cultures were induced with 1 mM IPTG and incubated overnight at 25 °C. Cells were pelleted and resuspended in GST lysis buffer [25 mM Tris-HCl (pH 7.5), 20 mM NaCl, 1 mM EDTA, 20% (v/v) glycerol, 1% (v/v) Triton X-100, and 2× protease inhibitor cocktail], then briefly sonicated (30-second pulses on/off for 5 minutes). Lysates were clarified by centrifugation at 13,000 rpm for 45 minutes. Protein concentrations were determined using BSA standards, and aliquots were stored at –80 °C.

Equimolar concentrations of GST-tagged proteins were immobilized onto glutathione-Sepharose beads by incubation in binding buffer [50 mM Tris-HCl (pH 7.4), 100 mM NaCl, 0.4% (v/v) Nonidet P-40, 10 mM MgCl₂, 5 mM EDTA, 2 mM DTT, and protease inhibitors] for 1 hour at 4 °C or overnight with gentle tumbling. After washing, beads were incubated with equimolar amounts of purified His-tagged wild-type or mutant GIV-CT proteins (resuspended in the same buffer) for 4 hours at 4 °C. Ten percent of the His-tagged input was retained as control.

Following incubation, beads were washed four times with phosphate wash buffer [4.3 mM Na₂HPO₄, 1.4 mM KH₂PO₄ (pH 7.4), 137 mM NaCl, 2.7 mM KCl, 0.1% (v/v) Tween-20, 10 mM MgCl₂, 5 mM EDTA, 2 mM DTT, 0.5 mM sodium orthovanadate], then eluted twice in Laemmli buffer [5% SDS, 156 mM Tris base, 25% glycerol, 0.025% bromophenol blue, 25% β-mercaptoethanol] by boiling at 95 °C for 5 minutes. Eluates were pooled, resolved by 10% SDS-PAGE, and transferred to PVDF membranes.

Membranes were stained with Ponceau S to confirm transfer efficiency and equimolar loading of GST-tagged proteins, then blocked in PBS containing 5% non-fat milk. Immunoblotting was performed using mouse anti-His antibody (Sigma; 1:1000 dilution) and a custom rabbit polyclonal anti-GIV-CT antibody (1:1000 dilution; epitope: C-terminal 18 amino acids of human GIV, validated previously^121^). Detection and quantification were carried out using a LI-COR Odyssey imaging system with dual-color infrared detection. Final figures were assembled using Adobe Photoshop and Illustrator (Adobe, San Jose, CA, USA).

### Alignment of Toll-interleukin-1 receptor (TIR) domain sequences

Amino acid sequences of the TIR-domain containing proteins were retrieved from the Ensembl Genome Browser (www.ensembl.org) and aligned using the Clustal Omega Multiple Sequence Alignment tool (https://www.ebi.ac.uk/Tools/msa/clustalo/). The resulting alignments were visualized using the BoxShade Server (https://embnet.vital-it.ch/software/BOX_form.html).

### Homology modeling of GIV•PD1 interaction via TIR-like BB-loop (TILL) sequences

In the absence of resolved structures of the cytoplasmic tail of PD1 and GIV’s C-terminus, we built homology models of the GIV(TILL)•PD1(TILL) complex using as template a previously published ^49,122^ model of GIV’s TILL motif bound to TLR4’s BB loop via a homotypic interface. Homology modeling was performed in ICM (Internal Coordinate Mechanics) software and the details of how the original GIV•TLR4(TIR) model was rigorously vetted and experimentally validated can be found in^49^. Briefly, the GIV-TILL and PD1-TILL peptides were built *ab initio*, tethered to the respective Pro-Gly positions, and its conformations were extensively sampled (> 108 steps) by biased probability Monte Carlo (BPMC) sampling in internal coordinates, with the TLR4 TIR domain represented as a set of energy potentials precalculated on a 0.5 Å 3D grid and including Van der Waals potential, electrostatic potential, hydrogen bonding potential, and surface energy. Following such grid-based docking, the peptide poses were merged with full-atom models of the TLR4 TIR domain, and further sampling was conducted for the peptide and surrounding side chains of the TLR4 residues. Co-complexed GIV-TILL and PD1-TILL peptide-only models are displayed here.

### Software and statistical analysis

All images were processed using ImageJ software, QuPath, Gen5, LAS X, or iStudio (LiCOR, Lincoln, NE, USA) software and assembled into figure panels with Photoshop and Illustrator (Adobe Creative Cloud).

All data are presented as mean ± standard error of the mean (SEM) from replicate experiments. Statistical analyses were performed using GraphPad Prism software (version 9.0). Comparisons between two groups were conducted using either Student’s *t*-test (parametric) or the Mann–Whitney *U*-test (non-parametric). For comparisons involving more than two groups, one-way or two-way analysis of variance (ANOVA) followed by Tukey’s post hoc test was used. A *p*-value of < 0.05 was considered statistically significant.

## DETAILED COMPUTATIONAL METHODS

### RNA Sequencing and Data Processing

RNA sequencing libraries were generated at the UCSD IGM Genomic Center using the Illumina TruSeq Stranded Total RNA Library Prep Gold with TruSeq Unique Dual Indexes (Illumina, San Diego, CA). Samples were processed following the manufacturer’s instructions, except for modifying the RNA shear time to five minutes. The resulting libraries were multiplexed and sequenced with 100 base pairs (bp) Paired-End (PE100) to a depth of approximately 40 million reads per sample on an Illumina NovaSeq 6000. Samples were demultiplexed using bcl2fastq v2.20 Conversion Software (Illumina, San Diego, CA).

Raw FASTQ files were trimmed, filtered, and mapped to the human genome for downstream quantification analyses. Low-quality sequences were trimmed or removed with Trimmomatic. STAR (version 2.6.0a) was used to align reads on the reference genome (*Mus musculus* genome build GRCm38.94). The resulting transcriptome-aligned sequences were used for expression quantification by using RSEM (version 1.3.3) with “–forward-prob 0” option. TPM scores for each sample were used through the RSEM tables (RSEM gene. results tables). We used log2(TPM) if TPM >1, else TPM – 1 values for each sample as the final gene expression value for downstream expression analyses.

Datasets are uploaded to NCBI GEO for both whole tumors (GSE314811) and TAM (GSE314813) datasets. Access codes to datasets are provided in “*Cover Letter*” and the datasets will be publicly released at the time of publication.

### Differential Expression Analysis

Differential gene expression analysis was performed using raw count data processed through DESeq2^123^. Genes with an absolute log2 fold change ≥ 1 and an adjusted p-value ≤ 0.05 (Benjamini-Hochberg method) were considered differentially expressed genes (DEGs). Pathway enrichment analysis of DEG lists was conducted using the ShinyGO ^124^ database and its integrated algorithms. Volcano plots representing differentially expressed genes were created using the matplotlib package.

### StepMiner analysis

StepMiner is an algorithm designed to detect stepwise transitions in time-series gene expression data^125^. It fits step functions to expression profiles by identifying the sharpest changes in signal, corresponding to gene expression switching events. The algorithm evaluates all possible step positions and calculates the average expression on either side of each step to define constant segments. An adaptive regression approach is then used to select the step position that minimizes the sum of squared errors between the observed and fitted data. The selected step is used as the *StepMiner* threshold*. This* threshold is used to convert gene expression values into Boolean values. A noise margin of 2-fold change is applied around the threshold to determine intermediate values, and these values are ignored during Boolean analysis. Finally, a regression test statistic is computed to assess the significance of the identified step transition as follows:

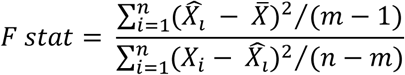

Where *X_i_* for *i* = 1 to *n* are the values, 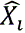 for *i* = 1 to *n* are fitted values. m is the degrees of freedom used for the adaptive regression analysis. 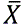 is the average of all the values: 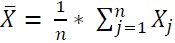. For a step position at k, the fitted values 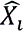 are computed by using 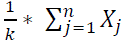 for *i* = 1 to *k* and 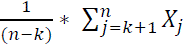 for *i* = *k* + 1 to *n*.

### Composite gene signature analysis using Boolean Network Explorer (BoNE)

Boolean network explorer (BoNE)^126^ provides an integrated platform for the construction, visualization and querying of a gene expression signature underlying a disease or a biological process in three steps: First, the expression levels of all genes in these datasets were converted to binary values (high or low) using the StepMiner algorithm. Second, Gene expression values were normalized according to a modified Z-score approach centered around *StepMiner* threshold (formula = (expr – SThr)/3*stddev). Third, the normalized expression values for every gene were added together to create the final composite score for the gene signature^127^. As a modified Z-score, the composite score of any gene signature typically ranges from negative to positive values, reflecting the dynamic range of each gene in the signature. These composite scores represent the overall activity or state of biological pathways associated with the genes and, hence, can identify differences between control and query groups within any given dataset. However, composite scores can neither be directly compared between various gene signatures on a dataset (because they are not normalized according to the number of genes in each signature), nor can the same signature be compared across datasets (which are individually normalized according to intra-dataset sample distribution or have other inherent differences). The samples were ordered based on the final signature score.

### Measurement of classification strength or prediction accuracy

To evaluate classification strength and prediction accuracy, Receiver Operating Characteristic (ROC) curves were generated for each gene. These curves assess the performance of a binary classifier, e.g., high vs. low *StepMiner*^125^ normalized gene expression levels, across varying discrimination thresholds. ROC curves plot the True Positive Rate (TPR) against the False Positive Rate (FPR) at multiple threshold levels. The Area Under the Curve (AUC) quantifies the classifier’s ability to correctly distinguish between the various types of samples, in this instance, WT vs KO tumors (GSE314811) or TAMs (GSE314813). ROC-AUC values were computed using the Python Scikit-learn package.

### Expression Heatmap

Z-score–normalized expression values of selected genes were visualized as a heatmap across samples within each dataset. Unsupervised clustering was performed to identify gene expression clusters using the Seaborn package (version 0.10.1).

### Bulk RNAseq Deconvolution

*In silico* deconvolution of bulk RNA-seq (tumor) data was performed using the Granulator R package^128^ to estimate immune cell compositions. Cell-type abundance estimates were normalized using the immune cell signature matrix developed by Monaco et al^129^.

### Survival Analysis

Kaplan–Meier survival analysis and log-rank tests for statistical significance were performed using KM-plotter (https://kmplot.com/analysis/)^130^. Briefly, datasets from immunotherapy treated patients with different tumor types including lung, melanoma, bladder cancer, head and neck, esophageal adenocarcinoma, hepatocellular cancer and glioblastoma were analyzed using the “Immunotherapy” subsystem^131^. The KM Plotter uses a default algorithm to automatically select the best cutoff value for separating samples into high and low expression groups when generating Kaplan-Meier survival plots. This automated selection is based on finding the cutoff that yields the most statistically significant separation between the groups, as determined by a Cox regression analysis^132^.

### Statistical Tests

Optimal gene expression cut-off values were determined using the *StepMiner* algorithm within each individual dataset^125^. Gene signatures were used to classify sample categories, and multi-class classification performance was evaluated using ROC-AUC (receiver operating characteristic area under the curve) values.

Violin and dot plots were generated using Seaborn (Python, version 0.12). Statistical comparisons were performed using Welch’s two-sample t-test (unpaired, unequal variance and sample size) via the scipy.stats.ttest_ind function in Python (version 0.19.0; equal_var=False). Multiple hypothesis correction was applied using the Benjamini-Hochberg method (fdr_bh) via statsmodels.stats.multitest.multipletests.

Differential expression analysis was conducted using DESeq2 in R (version 1.16.1).

## SUPPLEMENTAL INFORMATION INDEX

### Supplemental Figures and Legends (Figure S1-S7)

Figure S1: Functional annotation of responder and non-responder TAM gene programs, related to Figure 1.

Figure S2: Immune cell composition and spatial organization in the tumor microenvironment, related to Figure 2.

Figure S3: GIV levels dictate macrophage phagocytic efficacy and enable functional validation in patient-derived organoids, related to Figure 3.

Figure S4: Quantitative analysis of early tumor response to anti–PD1 therapy *in vivo*, related to Figure 4.

Figure S5: GIV-dependent tumor transcriptome does not predict clinical benefit from anti–CTLA-4 therapy, related to Figure 5.

Figure S6: GIV functions as an endocytic adaptor for PD1, related to Figure 6.

Figure S7: Somatic mutation landscape of PD1 in human cancers and structural impact on a regulatory surface patch affecting receptor positioning and function, related to Figure 7.

### Supplemental Tables (Table S1)

Table S1: Characteristics of patients enrolled into this study for obtaining lung tissues to serve as source of stem cells to generate lung organoids, related to Figure 3.

### Supplemental Data (Data S1-S5)

Data S1: Integration of SMaRT model with patient-derived signatures of ICI response, related to Figure 1.

Data S2: List of rTAM and nrTAM genes, related to Figure 1.

Data S3: Differentially expressed genes (DEGs) between WT and GIV-KO, whole tumors, related to Figure 5.

Data S4: Differentially expressed genes (DEGs) between WT and GIV-KO, F4/80+TAMs, related to Figure 5.

Data S5: Gene signatures of ICI-related immune responses and endocytic functions, related to Figure 5.

