## Supplemental details for "An Endocytic Checkpoint Controls Macrophage PD-1 Function and Immunotherapy Fate"

**AFFILIATIONS:**
[1]Department of Cellular and Molecular Medicine, University of California San Diego.
[2]Biomedical Sciences Graduate Program, School of Medicine, University of California San Diego, La Jolla, CA, USA.
[3]UC San Diego HUMANOID™ Center of Research Excellence, San Diego, CA, USA.
[4]Division of Cardiothoracic Surgery, University of California, 9300 Campus Point Drive, La Jolla, San Diego, CA 92037-7892, USA.
[5]Department of Medicine, University of California San Diego.

**Conflict of interest statement:** All authors have declared no conflicts of interest.

***Correspondence to**:

**Pradipta Ghosh, M.D.;** Professor, Departments of Medicine, and Cellular and Molecular Medicine, University of California San Diego; 9500 Gilman Drive (MC 0651), George E. Palade Building, Rm 232A, 239; La Jolla, CA 92093. Phone: 858-822-7633: Fax: 858-822-7636:

### CATALOG OF SUPPLEMENTAL INFORMATION

1. *Supplementary Figures and Legends (S1-S7)*
2. *Supplemental Tables (S1)*
3. *Supplementary Bibliography*

### SUPPLEMENTAL FIGURES AND LEGENDS

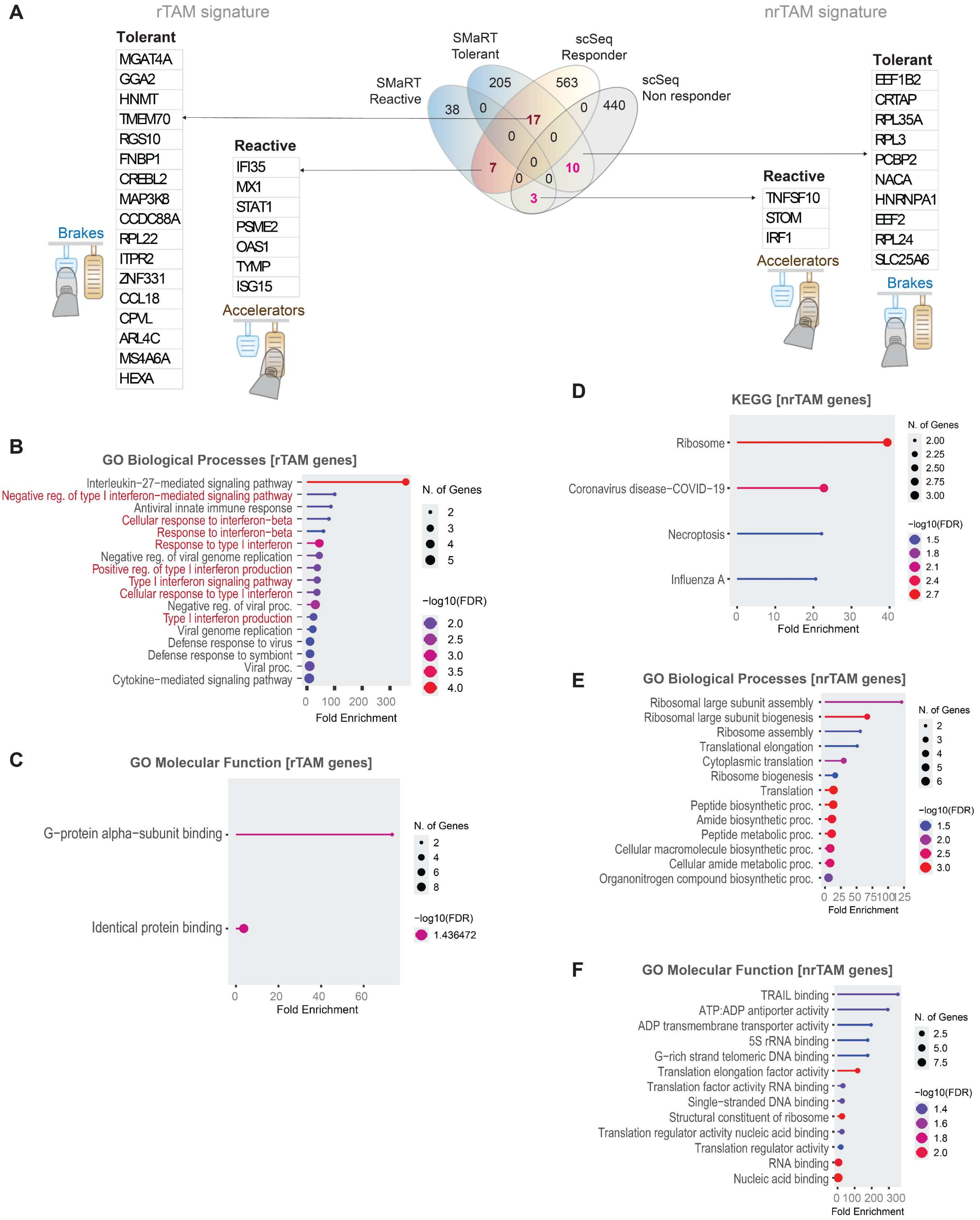


**Figure S1. [Related to Figure 1]**

**Functional annotation of responder and non-responder TAM gene programs.**

(**A**) Venn diagram showing overlap between a computational macrophage continuum model—Signatures of Macrophage Reactivity and Tolerance (SMaRT)[1] and pan-cancer single-cell–derived tumor-associated macrophage (TAM) gene sets stratified by clinical response to immune checkpoint blockade (responder, rTAM; non-responder, nrTAM)[2]. The SMaRT framework captures invariant macrophage states across >12,500 transcriptomic datasets, organized into reactive "accelerator" modules and tolerant "brake" modules.
(**B-C**) Gene Ontology (GO) enrichment analyses of rTAM genes overlapping the SMaRT model, highlighting biological processes and molecular functions associated with antiviral defense, interferon signaling, and G protein–related activities.
(**D**) KEGG pathway enrichment of nrTAM genes, revealing pathways linked to translational control, ribosomal function, and stress-associated programs.
(**E-F**) GO biological process and molecular function enrichment analyses of nrTAM genes, indicating dominance of biosynthetic, translational, and RNA-binding activities characteristic of macrophage states associated with therapeutic non-response.

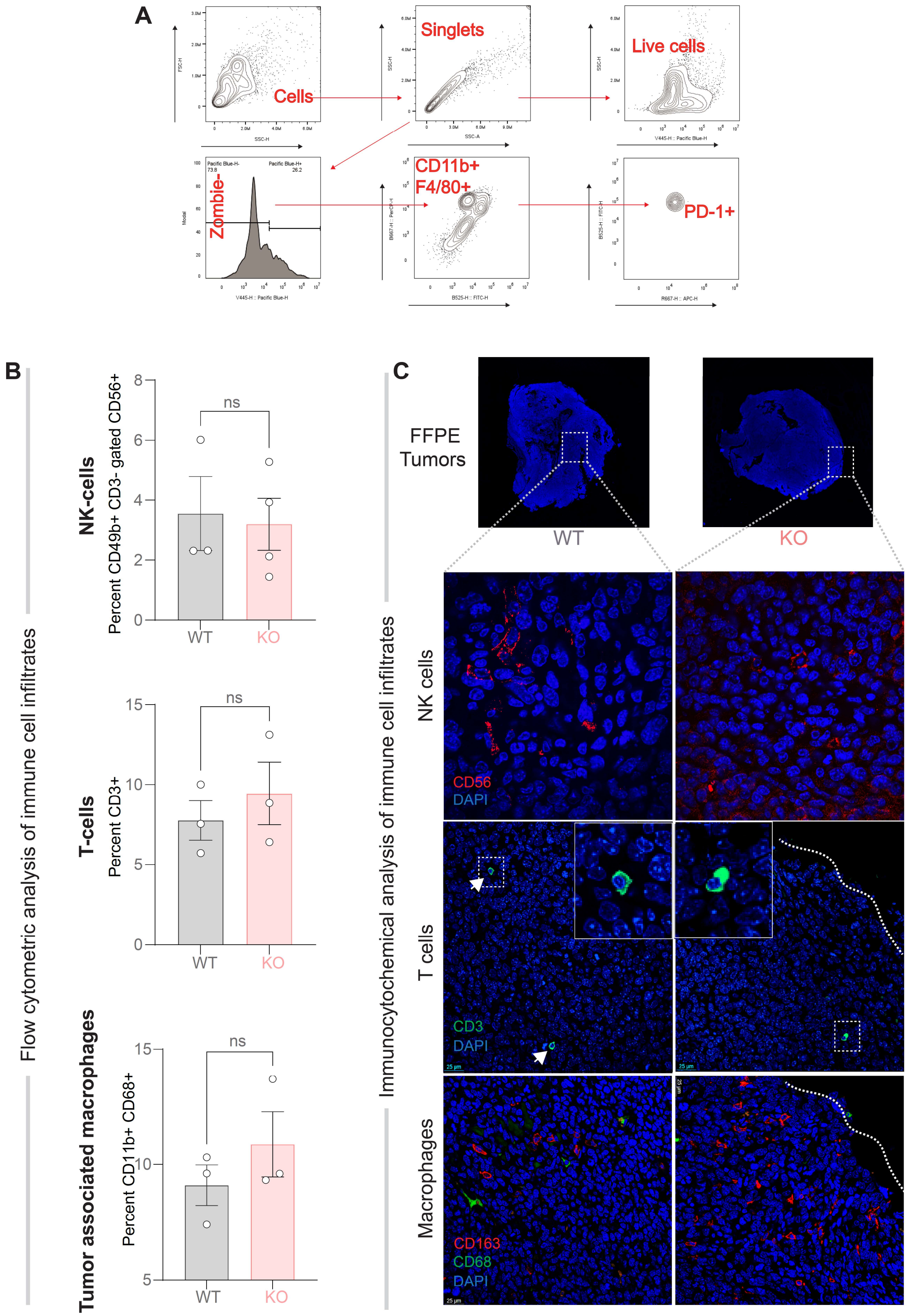


**Figure S2. [Related to Figure 2]**

**Immune cell composition and spatial organization in the tumor microenvironment.**

(**A**) Flow cytometry gating strategy used to identify tumor-associated macrophages (TAMs) from enzymatically dissociated tumor tissues. Sequential gates were applied to select total cells, singlets, live (Zombie$^{-}$) cells, and myeloid populations defined as CD11b$^{+}$F4/80$^{+}$, followed by assessment of surface PD1 expression.

(**B**) Flow cytometry-based assessment of relative abundance of immune cells in tumors from WT and GIV-KO mice.

(**C**) Spatial characterization of immune infiltrates in FFPE tumor sections. (Top) Whole-scan confocal images of representative WT and CCDC88A-KO tumors showing nuclear staining with DAPI (blue) and regions selected for highermagnification analysis. (Bottom) Confocal images illustrating immune cell subsets within the tumor microenvironment, including NK cells (CD56, red), T cells (CD3, green), and macrophages (CD68, green; CD163, red), with nuclei counterstained by DAPI (blue). Dashed outlines demarcate tumor boundaries and insets highlight representative immune cell localization within tumor cores and peripheries.

*Statistics:* Data are presented as mean ± SEM. Statistical significance was assessed using unpaired t tests, as indicated. $p \leq 0.05$ was considered significant.

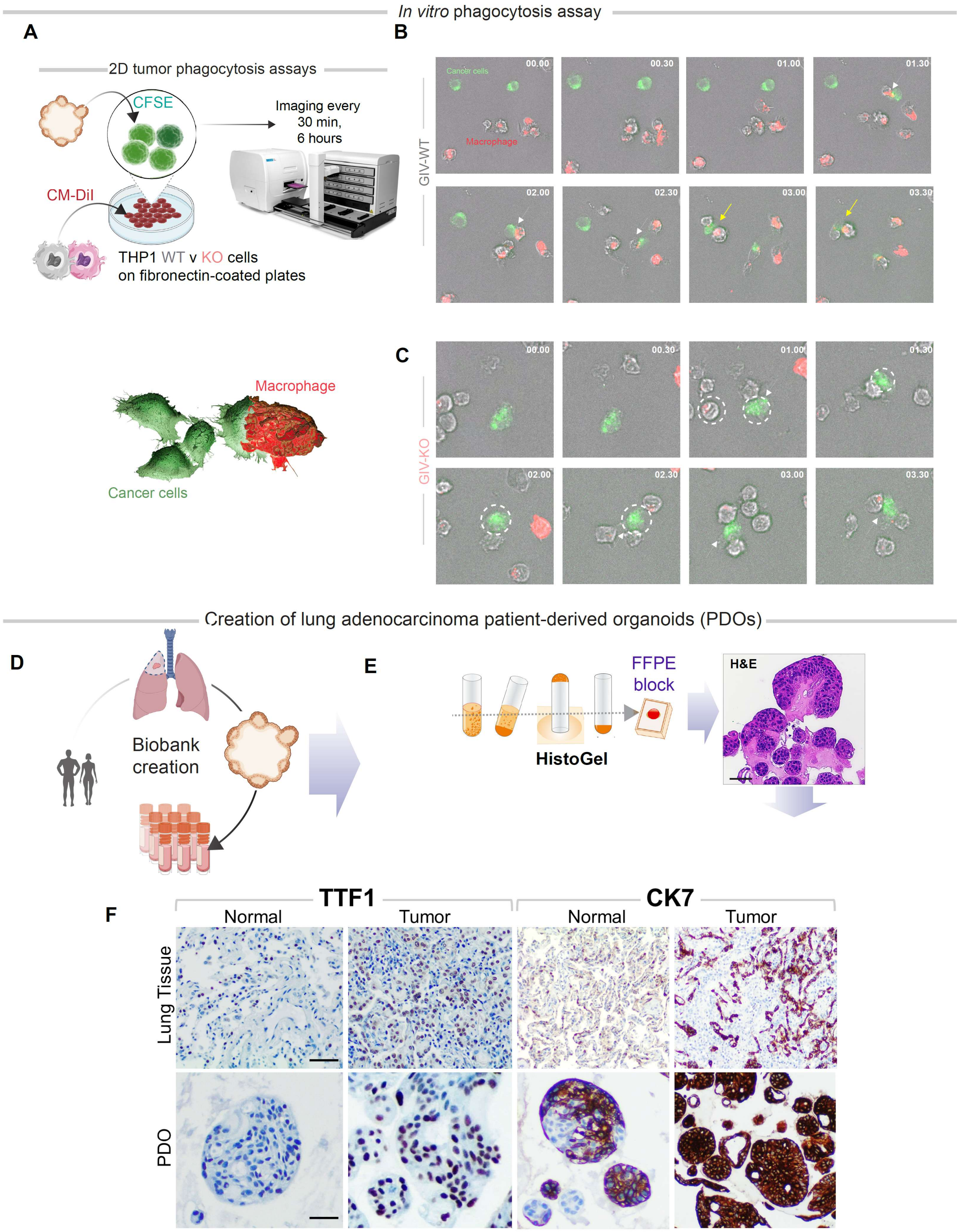


**Figure S3. [Related to Figure 3]**

**GIV levels dictate macrophage phagocytic efficacy and enable functional validation in patient-derived organoids.**

(**A**) Schematic of the 2D in vitro phagocytosis assay used to assess macrophage-mediated tumor cell uptake. PMA-differentiated THP1 macrophages (WT or GIV-KO) were labeled with CM-DiI (red) and co-cultured on fibronectin-coated plates with CFSE-labeled MDA-MB-231 cancer cells (green), followed by time-lapse imaging every 30 min for up to 4 h.
(**B**-**C**) Representative time-stamped confocal images obtained using Cytation 10 (Agilent) illustrating dynamic interactions between cancer cells (green) and THP1-derived macrophages (red). Arrows in panel B track multiple serial frames that capture GIV-proficient WT macrophages engulfing a tumor cell. Interrupted circles in panel C track multiple serial frames in which a single tumor cell survive multiple encounters with numerous GIV-deficient macrophages.
(**D**) Workflow for the generation of patient-derived lung adenocarcinoma organoids (PDO-LUAD), including tissue sampling, isolation, and biobank establishment.
(**E**) Schematic of histogel embedding, FFPE block preparation, and hematoxylin and eosin (H&E) staining of PDOs for histopathologic validation.
(**F**) Immunohistochemical analysis of lung adenocarcinoma markers TTF1 (left) and CK7 (right) in matched primary lung tissues (top) and corresponding PDOs (bottom). Normal lung PDOs exhibit intra- and inter-organoid staining heterogeneity, whereas tumor-derived PDOs show strong, homogeneous marker expression. Scale bar, 50 μm. Representative images from five independent PDO isolations (100% success rate) are shown.

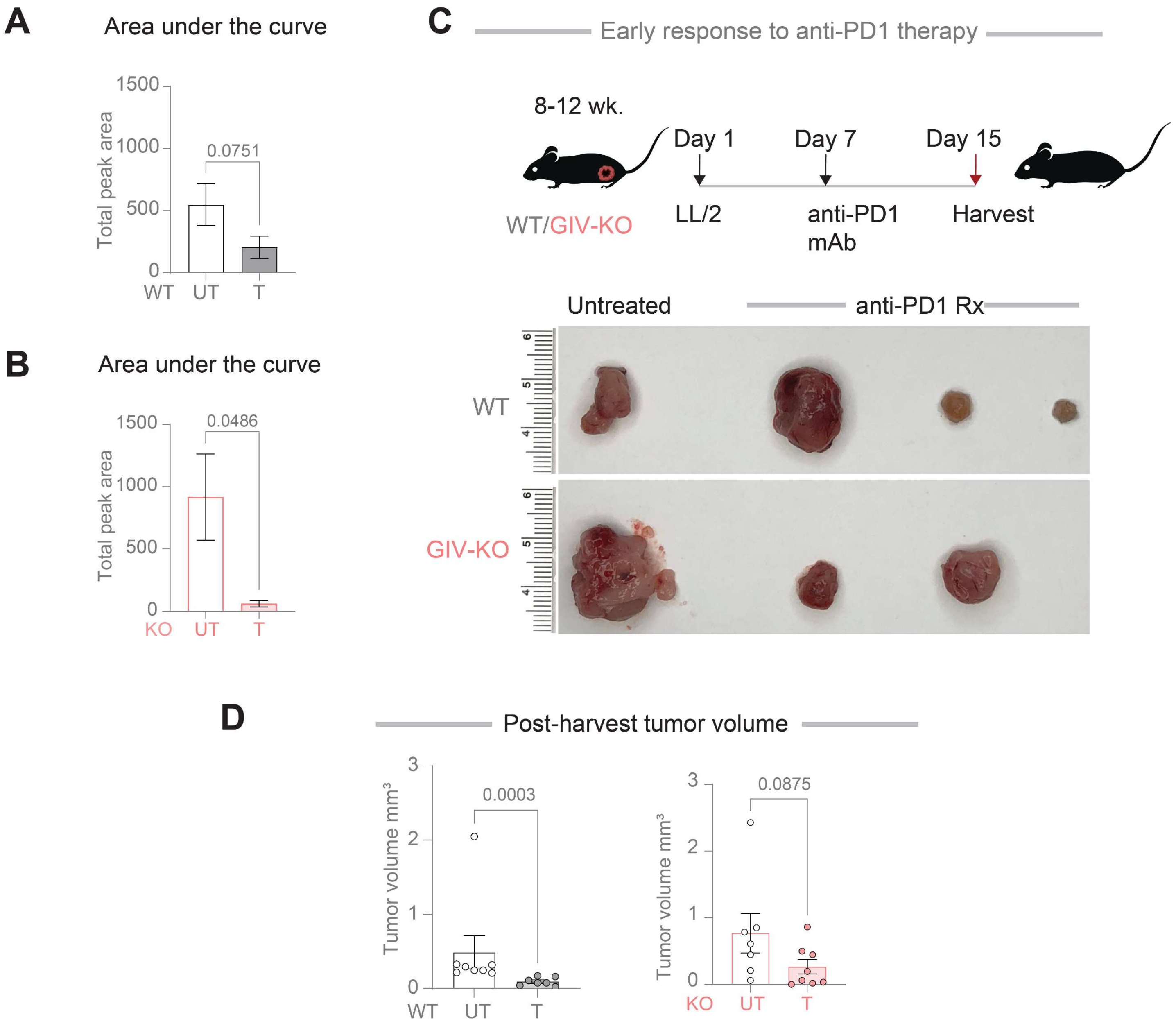


**Figure S4. [Related to Figure 4]**
**Quantitative analysis of early tumor response to anti–PD1 therapy *in vivo*.**
(**A-B**) Area-under-the-curve (AUC) analyses of longitudinal tumor growth in response to anti–PD1 monoclonal antibody treatment in WT mice (A) and myeloid-specific GIV-KO mice (B), highlighting comparable early therapeutic responses between genotypes.
(**C**) Experimental schematic for assessing early response to anti–PD1 therapy. WT and GIV-KO mice bearing LL/2 tumors were treated with anti–PD1 beginning on day 7 after tumor implantation and tumors were harvested at day 15. Representative post-harvest tumor images from untreated and anti–PD1–treated mice are shown.
(**D**) Quantification of post-harvest tumor volumes at the 15-day time point in WT and *CCDC88A*-KO mice, demonstrating similar early tumor control in WT and GIV-deficient mice.

*Statistics:* Data are presented as mean ± SEM. Statistical significance was assessed using unpaired t tests, as indicated. $p \leq 0.05$ was considered significant.

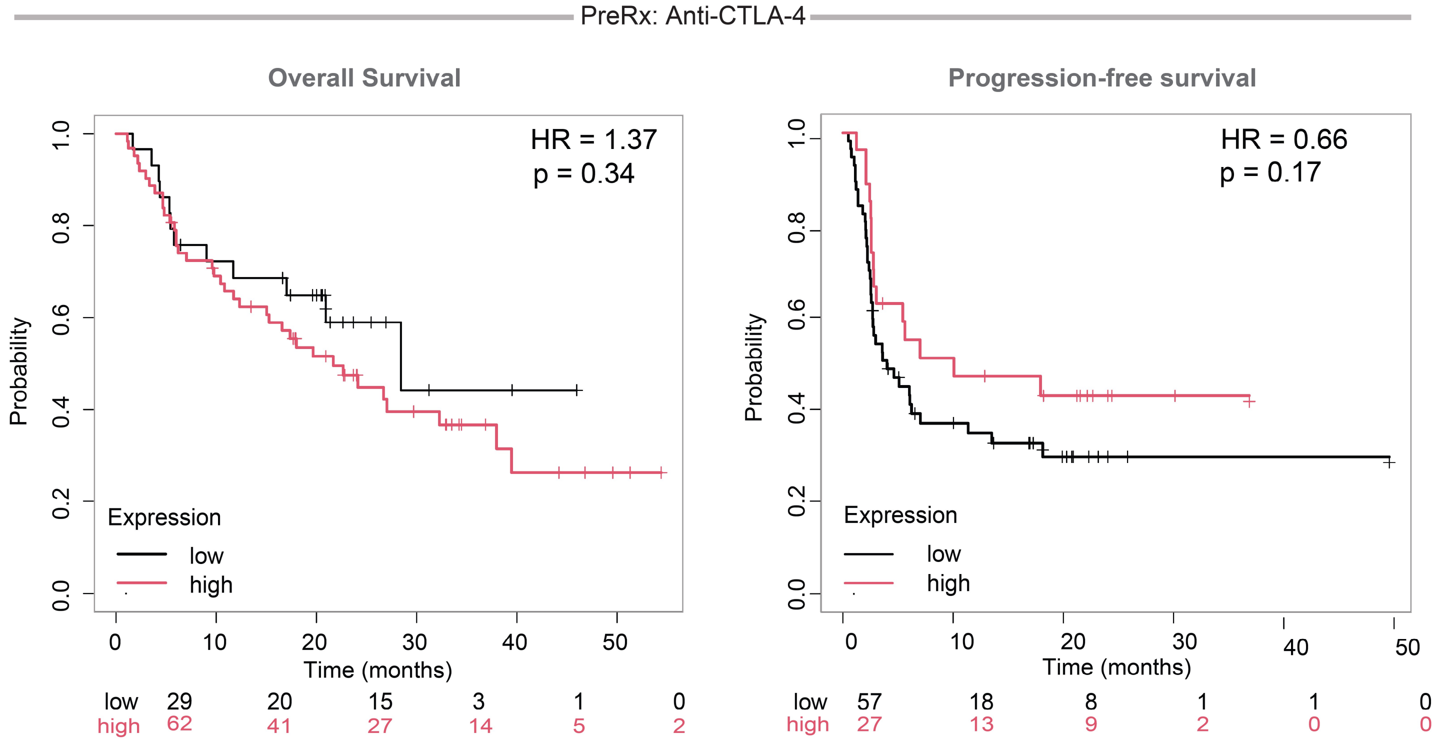


**Figure S5. [Related to Figure 5]**

**GIV-dependent tumor transcriptome does not predict clinical benefit from anti–CTLA-4 therapy.**

Kaplan–Meier analyses showing the association between expression of the GIV-linked tumor gene signature—identified as upregulated in tumors from *CCDC88A* WT mice—and clinical outcomes in patients treated with anti–CTLA-4 therapy. Overall survival (left) and progression-free survival (right) are shown for patients stratified by high versus low expression of the signature. In contrast to PD1/PD-L1–based immunotherapy (**Figure 5E**), GIV-associated gene expression does not significantly predict response to CTLA-4 blockade, underscoring the pathway specificity of GIV-dependent macrophage PD1 trafficking in shaping therapeutic outcomes.

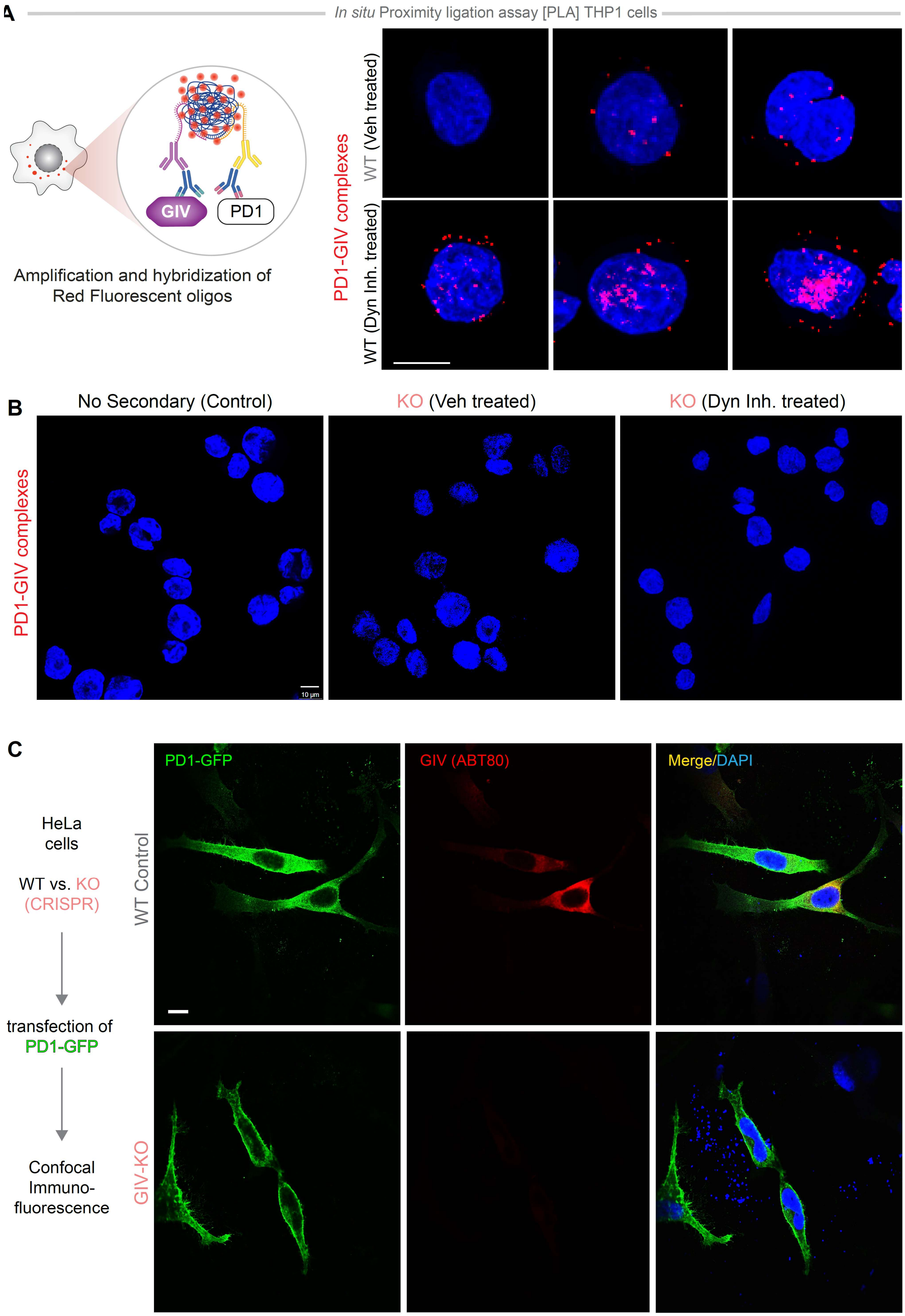


**Figure S6. [Related to Figure 6]**

**GIV functions as an endocytic adaptor for PD1.**

(**A**) In situ proximity ligation assay (PLA) used to detect endogenous PD1–GIV complexes in THP1 macrophages. (Left) Schematic illustrating PLA principle, in which proximal binding of antibodies recognizing PD1 and GIV enables amplification and hybridization of fluorescent oligonucleotides, visualized as discrete red puncta. (Right) Representative confocal images showing PD1–GIV PLA signals (red) in nuclei counterstained with DAPI (blue). Increased puncta are observed upon inhibition of dynamin-dependent endocytosis, consistent with stabilization of PD1–GIV complexes at endomembranes.

(**B**) Specificity controls for PLA. Confocal images from "no secondary antibody" controls and from vehicle-treated versus DynGo4a-treated THP1 cells demonstrate absence of nonspecific PLA signals, confirming assay specificity for PD1–GIV interactions.

(**C**) Validation of PD1–GIV association in a heterologous system. (Left) Experimental schematic showing CRISPR-engineered WT or GIV-KO HeLa cells (extensively validated previously)[3-5] transiently transfected with GFP-tagged PD1. (Right) Confocal immunofluorescence images showing PD1–GFP (green), endogenous GIV (red), and nuclei (DAPI, blue). Co-localization of PD1 and GIV is evident in WT cells and is lost upon GIV deletion, supporting a direct role for GIV in PD1 trafficking.

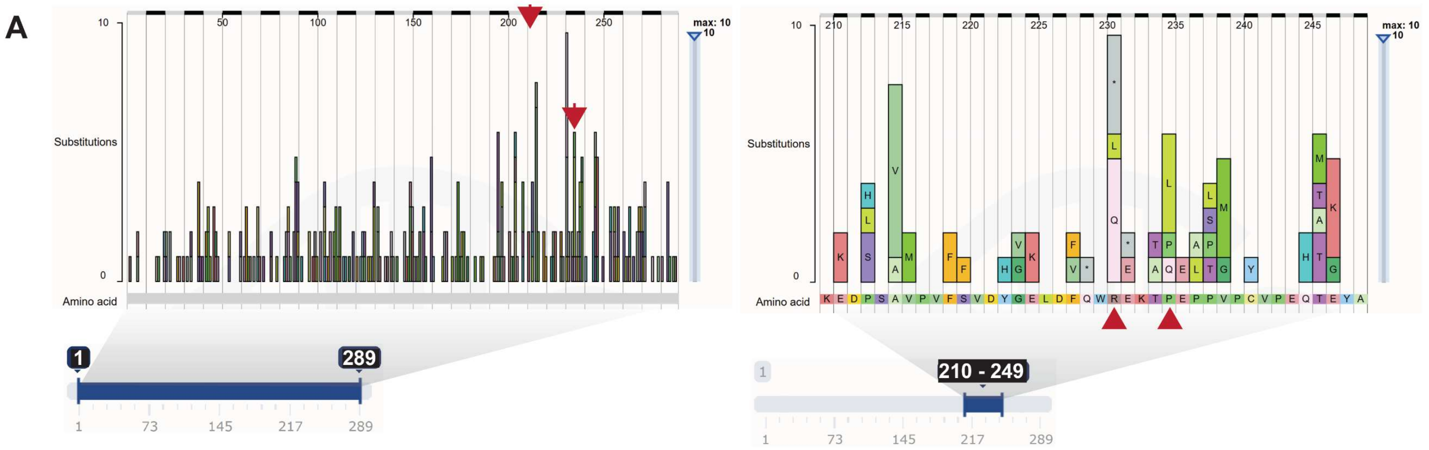


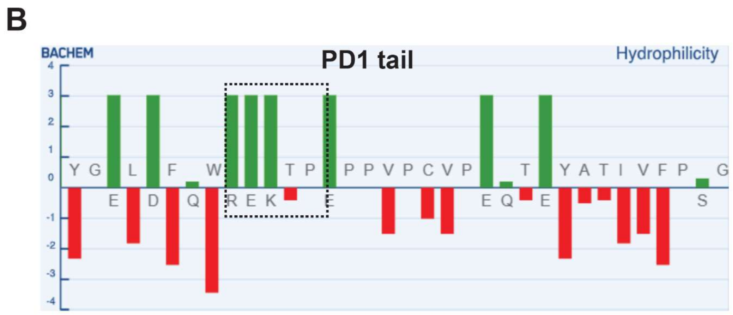


| Sequence | Isoelectric point | Net charge at pH 7.0 |
|---|---|---|
| Entire tail | 5.79 | -1.03 |
| REKTP | 10.12 | -1.03 |
| QEKTP | 6.97 | -1.00 |
| LEKTP | 6.97 | 0.00 |
| REKTQ | 10.12 | 1.00 |
| REKTL | 10.12 | 1.00 |

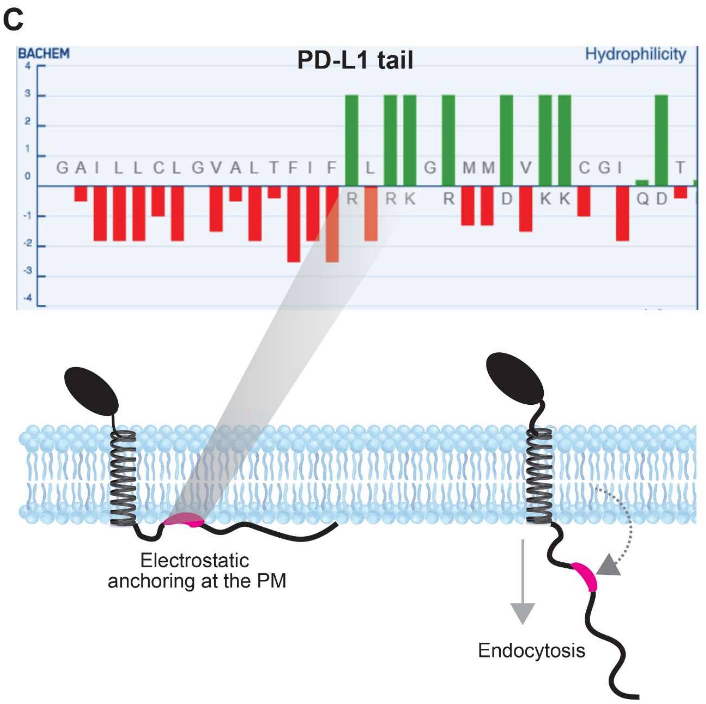


**Figure S7**. [**Related to Figure 7**]

**Somatic mutation landscape of PD1 in human cancers and structural impact on a regulatory surface patch affecting receptor positioning and function**.

(**A**) Lollipop plot summarizing somatic mutations in the PD1 gene across diverse cancer types from the Catalogue Of Somatic Mutations In Cancer (COSMIC), colored by mutation class (missense, nonsense, frameshift, splice). The PD1 topography (left) across tumors shows clustering of recurrent substitutions within discrete protein segments, including the extracellular IgV domain and juxtamembrane region, suggesting potential hotspots for altered receptor stability or ligand engagement (COSMIC overview). Zoomed-in view (right) within the cytosolic domain highlights a membrane proximal region (aa 210-249) within which arrowheads (red) indicate the substitutions that impair PD1 interaction with GIV.

(**B**) Hydrophilicity plot (Y-Axis, hydropathy index; X-Axis, amino acid sequence number) of the boxed sequence within PD1 tail, and the impact of substitutions on the charge. Positive Peaks: Indicate hydrophobic stretches (e.g., membrane-spanning alpha-helices). Negative Troughs: Indicate hydrophilic regions (e.g., surface-exposed parts in water-soluble proteins).

(**C**) Proposed mechanism whereby somatic alterations within this patch in panel (B) perturb PD1 positioning on the plasma membrane or its stability, potentially phenocopying mechanisms reported previously for a similar 'hydrophobic patch' in PD-L1 cytosolic tail [6], where specific loops and short sequence patches govern localization, endocytic recycling, and immune evasion in tumors.

Together, these data integrate cancer mutation cataloging with experimental insights, suggesting that selected PD1 somatic mutations cluster in regions critical for receptor endocytosis, positioning and function on the cell surface, with implications for immune checkpoint modulation and therapeutic sensitivity in oncology.

### SUPPLEMENTAL TABLES

**Table S1 [Related to Figure 3]. Characteristics of patients enrolled into this study for obtaining lung tissues to serve as source of stem cells to generate lung organoids.**

| Name | Tumor collected | Age | Sex | Smoking history | Reason for surgery | Histology |
|---|---|---|---|---|---|---|
| **JA-5** | Yes | 46 | Female | Non-smoker | Left lower lobe nodule | Invasive adenocarcinoma and adjacent normal |
| **JA-14** | Yes | 74 | Female | Former Smoker 1 pack/day Duration:40 years | Left upper lobe mass | Adenocarcinoma and adjacent normal |
